# Maternal Folate Nutrition Programs Adult Hematopoietic Stem Cell Function Through Mitochondrial Remodeling

**DOI:** 10.64898/2026.10.26.761955

**Authors:** Brian P. Krum, Ava Niazi, Victoria Chiou, Brittany M. Curtiss, Noah Huerta, Fanny Zhou, Luisa F. Castillo, Sweta B. Patel, Alana M. Franceski, Sajesan Aryal, Anton Classen, Travis Nemkov, Angelo D’Alessandro, Martha S. Field, Rui Lu, Robert A.J. Signer, Robert S. Welner, Anna E. Beaudin

## Abstract

Folic acid fortification for the prevention of neural tube defects has been a cornerstone of public health policy for nearly three decades, but its broader consequences for other developing organ systems, particularly the hematopoietic system, are poorly studied. Hematopoietic stem cell (HSC) emergence overlaps developmentally with neural tube closure, and folate-dependent one-carbon metabolism is critical for nucleotide synthesis, cellular methylation, and redox balance, positioning folate as a candidate regulator of HSC programming. Using dietary models of maternal folate deficiency or supplementation in mice, we demonstrate that maternal folate acts as a dose-dependent developmental tuner of hematopoiesis, programming adult offspring HSC function through divergent mitochondrial and chromatin states. Maternal folate deficiency impaired adult HSC self-renewal, associated with mitochondrial defects, disrupted cristae, elevated proteotoxic stress and a stress-adapted chromatin landscape. In contrast, maternal folate supplementation enhanced HSC self-renewal accompanied by improved mitochondrial quality control and CDC42-associated organelle remodeling, and established chromatin states permissive for stem cell maintenance. Fetal HSC metabolomics identified shifts in one-carbon intermediates, pyruvate and lipid pools, suggesting that divergent fetal metabolic states provide substrate-level inputs that drive persistent and opposing chromatin programs in offspring HSCs. Our findings establish maternal folate availability as an unappreciated regulator of lifelong hematopoietic output demonstrating that a single maternal micronutrient, folate, is sufficient to program adult HSC function and regenerative capacity through persistent metabolic and mitochondrial programming.

## INTRODUCTION

Population-wide dietary fortification of folic acid was implemented in 1998 in the United States after folic acid supplementation was shown to reduce neural tube defects (NTD) incidence by approximately 70%.^1–4^ Pregnant women and women planning pregnancies are also advised to take additional prenatal folate supplements. Although frank folate deficiency is relatively uncommon in many populations, suboptimal folate status remains prevalent among women of reproductive age, with a substantial proportion falling below the 906 nmol/L red blood cell folate concentration associated with optimal NTD prevention, including in folate-fortified populations such as the United States.^5,6^ Furthermore, additional evidence remains for genetically susceptible populations due to common polymorphisms in folate-mediated one carbon metabolism.^7–10^ Emerging evidence also raises questions about the safety of current fortification and supplementation practices, as excess folate intake can result in circulating unmetabolized folic acid.^11–15^ Although folic acid supplementation represents an unprecedented public health success story, the precise mechanism by which folic acid supplementation prevents NTDs is still elusive. Furthermore, the effects of prenatal folate nutrition on developing stem cells and other organ systems are largely unexplored.

Hematopoietic stem cell (HSC) emergence occurs approximately coincident with neural tube closure in both mice and humans.^16–18^ HSC development and function is regulated by outputs of folate-mediated one-carbon metabolism (OCM), including nucleotide synthesis, cellular methylation, and redox balance, ^19–22^ positioning the hematopoietic system to translate early-life nutritional cues into programmed outcomes on offspring biological circuits, with persistent effects on cellular metabolism and epigenetic state beyond the period of exposure.^23,24^ Nucleotide biosynthesis modulates HSC proliferation and genome maintenance.^25^ Epigenomic maintenance is a master regulator of HSC development and self-renewal^26^, and dysregulation is a risk factor for clonal expansion and leukemic transformation.^27,28^ Metabolic regulation is a central determinant of HSC fate and longevity;^29–31^ HSCs maintain a tightly controlled mitochondrial state,^32–35^ and disruption of mitochondrial metabolism impairs HSC fitness.^34,36–39^ Folate-mediated OCM is therefore strongly positioned to regulate multiple cellular outcomes involved in programming HSC function during development.

Whether maternal folate nutrition has lasting consequences for HSC function remains an important unresolved question. Here, we tested the effect of varying maternal folate nutrition on HSC function in adult offspring. Using established dietary models of maternal folate deficiency and supplementation,^40–43^ we show that maternal folate nutrition durably alters metabolic state and functional capacity of adult HSCs. Folate deficiency impaired self-renewal and mitochondrial integrity in adult offspring HSCs, whereas supplementation enhanced HSC regenerative potential. Enhanced self-renewal in supplemented HSCs was associated with transcriptional programs linked to mitochondrial quality control, autophagy, and CDC42-dependent cytoskeletal signaling. ATAC-seq profiling revealed divergent chromatin accessibility at metabolic and self-renewal loci, supporting a model in which folate-driven metabolic shifts in fetal HSCs are translated into lasting chromatin states that perpetuate altered HSC function. Our findings establish a new paradigm wherein maternal nutrient availability can permanently encode metabolic state in adult HSCs with persistent functional outcomes.

## METHODS

### Mice and dietary interventions

C57BL/6J, Flk2-Cre143^44^; mT/mG ^45^ (FlkSwitch), and congenic CD45.1 PepBoyJ mice were maintained under standard conditions. Additional details can be found in the supplemental Methods.

### Flow cytometry

Bone marrow cells were isolated as previously described.^46^ Additional details can be found in the supplemental Methods.

### Metabolomics analysis

Untargeted metabolomics was performed on sorted HSCs from three mice per biological replicate (n = 3/group). Metabolites were analyzed by UHPLC-MS (Vanquish UHPLC/Q Exactive; Thermo Fisher Scientific) as described.^47^ Additional details can be found in the supplemental Methods.

### Imaging and microscopy Transmission electron microscopy

A previously identified protocol was used.^48^ Additional details can be found in the supplemental Methods.

### Extracellular flux analysis

Mitochondrial respiration and glycolytic function were assessed in sorted fetal HSPCs using Seahorse XF Mito Stress and Glycolysis Stress Tests (Agilent Technologies).

### Immunocytochemistry and Super-resolution confocal imaging and 3D reconstruction

Sorted HSCs were fixed, permeabilized, and stained for TOMM20, CDC42, and LAMP1. Additional details can be found in the supplemental Methods.

### Genomic Analysis

Single-cell RNA-seq of Sorted adult offspring HSPCs was performed using the Chromium Single Cell 3′ platform (10x Genomics) and Illumina NovaSeq. Annotation using established hematopoietic markers.^49,50^ ATAC-seq data were processed using nf-core/atacseq ^51^ and differential accessibility was assessed using DESeq2.^52^ Additional details can be found in the supplemental Methods

### Total Folate Measurements

Plasma and tissue folate concentrations were quantified by Lactobacillus casei microbiological assay and normalized to protein as described.^53,54^

## RESULTS

### Maternal folate nutrition drives persistent alterations to adult offspring HSC function

To examine how maternal folate status shapes adult hematopoiesis, pregnant dams were maintained on folate deficient (Def, 0 mg/kg folic acid), control (2 mg/kg), or supplemented (Supp, 8 mg/kg) diets (**Table S1**), modeling human folate intake. Offspring were weaned onto standard chow, and adult bone marrow (BM) hematopoiesis was assessed (**Fig. 1A**). Maternal serum folate levels differed across diets (**Fig. S1A**), whereas total fetal liver (FL) folate at embryonic day (E)14.5 was unchanged (**Fig. S1B**). Dietary folate manipulation did not induce anemia or alter white blood cell counts in dams or neonatal offspring (**Fig. S1C–M**). Intracellular folate levels were modestly reduced in folate Def FL long-term (LT)-HSCs but unchanged in Supp LT-HSCs (**Fig. S1N**). By adulthood, intracellular folate levels in BM LT-HSCs were comparable across all diet groups (**Fig. S1O**).

**Figure 1.**
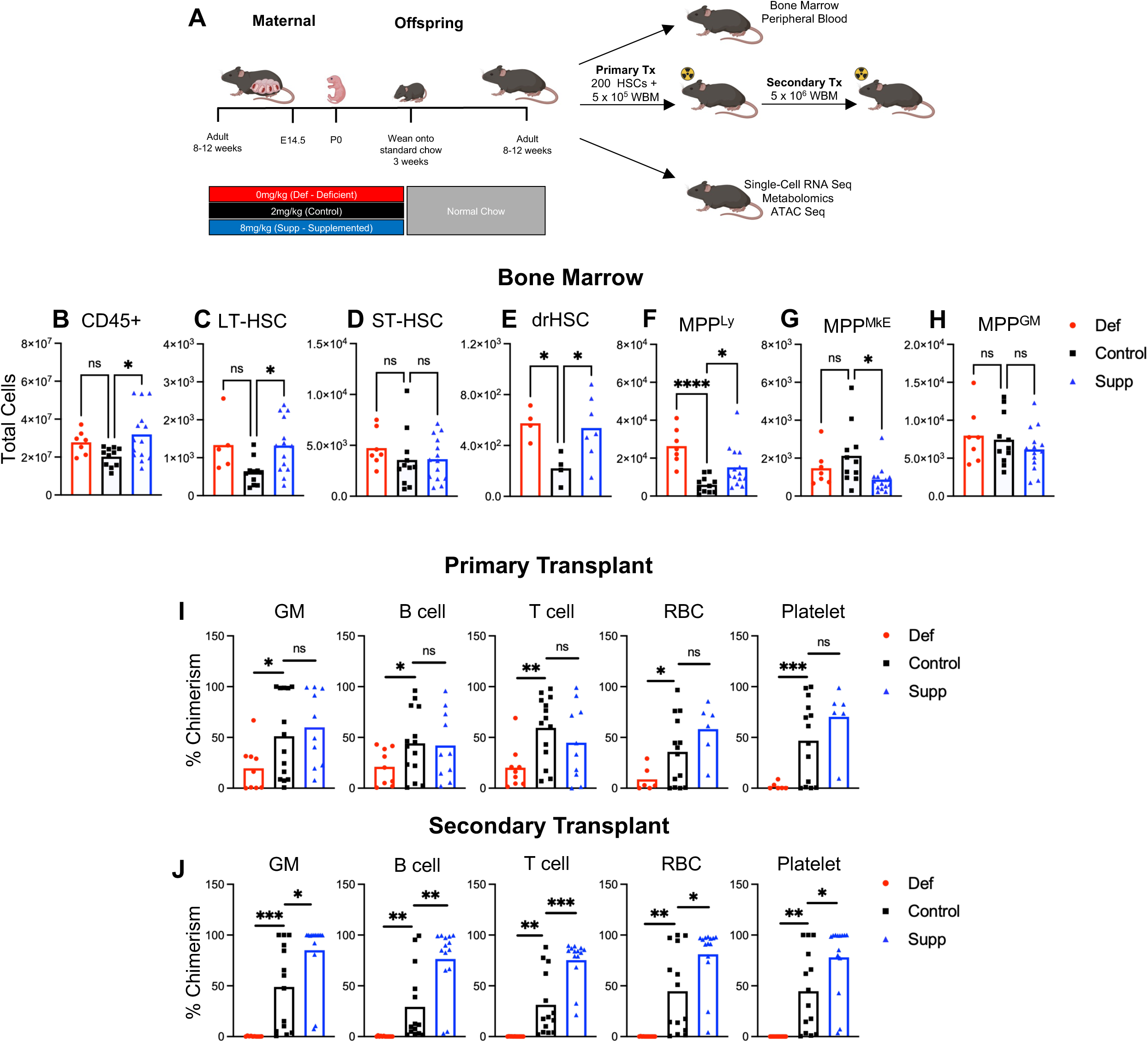
Maternal folate nutrition drives persistent alterations to adult offspring HSC function. **A)** Representative schematic of prenatal folate dietary intervention and experimental assessment of offspring. **B-H)** Total cellularity of bone marrow hematopoietic cell populations quantified by flow cytometry. (**B**) CD45^+^ cells, (**C**) LT-HSC, (**D**) ST-HSC, (**E**) Developmentally-restricted HSC (drHSC), (**F**) MPP4^Ly^ (**G**) MPP2^MkE^, (**H**) MPP3^GM^. **I)** Peripheral blood chimerism in at 16 weeks post-transplant in Granulocyte/Macrophage (GM), B-cell, T-cell, Red blood cells (RBC), and platelet lineages in primary recipients of 200 HSCs isolated from Def and Supp adult offspring bone marrow. **J)** Peripheral blood chimerism at 16 weeks post-transplant in GM, B-cell, T-cell, RBC and platelet lineages in secondary whole bone marrow recipients of Def and Supp adult offspring HSC recipients. For all experiments, N = 3 independent experiments representing three independent litters per group. Significance determined by one-way ANOVA with multiple comparisons or unpaired student’s t-test; *P < 0.05; **P < 0.01; ***P < 0.001; ****P<0.001.

Profiling of adult BM hematopoiesis revealed persistent effects of maternal folate intake (**Fig. 1**). Total CD45⁺ BM cellularity was increased in Supp offspring (**Fig. 1B**), accompanied by expansion of LT-HSCs (**Fig. 1C**) but not short-term (ST)-HSCs (**Fig. 1D**). Lymphoid-biased developmentally restricted HSCs (drHSCs) ^55^ and MPP4^Ly^ progenitors, previously shown to expand in response to prenatal inflammation,^46^ were expanded in both Def and Supp offspring (**Fig. 1E–F**), as were common lymphoid progenitors (CLPs; **Fig. S2A**). However, Def offspring showed no increase in mature BM B or T cells (**Fig. S2B–C**), whereas Supp offspring exhibited increased BM T cells (**Fig. S2C**). Megakaryocyte progenitors (MkP) were increased in Supp offspring (**Fig. S2D**) despite reduced erythroid-biased MPP2^MkE^ (**Fig. 1G**). Granulocytes were increased in both Def and Supp offspring (**Fig. S2E**), although myeloid-biased MPP3^GM^ progenitors were unchanged (**Fig. 1H**). Most changes were not reflected in peripheral blood (PB; **Fig. S2F–G**), except for reduced T cells in both Def and Supp offspring (**Fig. S2H**). Together, these findings demonstrate that maternal folate status programs persistent changes to adult BM hematopoiesis.

To test cell-autonomous alterations in LT-HSCs, we performed adoptive transfer assays with BM LT-HSCs from adult offspring (**Fig. 1A**). Def HSCs exhibited markedly impaired reconstitution across mature lineages over 16 weeks, whereas Supp HSCs maintained overall donor chimerism (**Fig. 1I**). BM analysis confirmed reduced reconstitution of most HSPC populations in Def HSC recipients (**Fig. S2I-J**). We next assessed long-term self-renewal potential in a secondary transplantation assay. Def HSCs failed to reconstitute recipients, whereas Supp HSCs exhibited pronounced enhancement of self-renewal with enhanced multilineage chimerism (**Fig. 1J**). Thus, maternal folate status is sufficient to program adult HSC function, altering both BM hematopoiesis and long-term HSC self-renewal capacity.

### Maternal folate deficiency imprints a dysfunctional mitochondrial metabolic program in offspring HSCs

To define metabolic changes associated with HSC programming by maternal folate, we performed untargeted mass spectrometry-based metabolomics on adult offspring HSCs. Partial least squares discriminant analysis and hierarchical clustering of the top 50 variable metabolites separated Def and Control HSCs (**Fig. S3A-B**), revealing 35 differentially abundant metabolites in Def HSCs compared with only 5 in Supp HSCs (**Fig. S3C-D**). Pathway analysis identified purine metabolism, glycolysis, and aminoacyl-tRNA biosynthesis as the most altered pathways in Def HSCs (**Fig. 2A**). Def HSCs exhibited increased glucose and lactate with reduced glucose-6-phosphate (G6P) and pyruvate levels (**Fig. 2B**), consistent with increased glycolytic flux and altered mitochondrial pyruvate utilization. Additional signatures indicated impaired oxidative metabolism, including an increased succinate:fumarate ratio consistent with Complex II dysfunction ^56^ (**Fig. 2C**) and accumulation of short- and medium-chain acylcarnitines, suggesting impaired β-oxidation ^57,58^ (**Fig. 2D**). Def HSCs also showed increased glutamine levels (**Fig. 2E**), elevated ATP, reduced ADP (**Fig. 2F-G**), and increased uracil abundance (**Fig. 2H**), consistent with altered nucleotide metabolism.^59^ Aminoacyl-tRNA biosynthesis enrichment was reflected in depletion of arginine, histidine, and serine (**Fig. 2I**), indicating remodeling of amino acid utilization and translational capacity. Together, these findings demonstrate that maternal folate deficiency profoundly remodels adult HSC metabolism, driving an increased reliance on glycolysis, mitochondrial dysfunction, amino acid metabolism and disrupted nucleotide homeostasis. We subsequently examined mitochondrial morphology by transmission electron microscopy (TEM). Def HSCs displayed abnormal mitochondrial ultrastructure, reduced and disorganized cristae and increased inner mitochondrial space (**Fig. 2J–K; Fig. S3E**), consistent with impaired mitochondrial function identified by metabolomics.^60,61^

**Figure 2.**
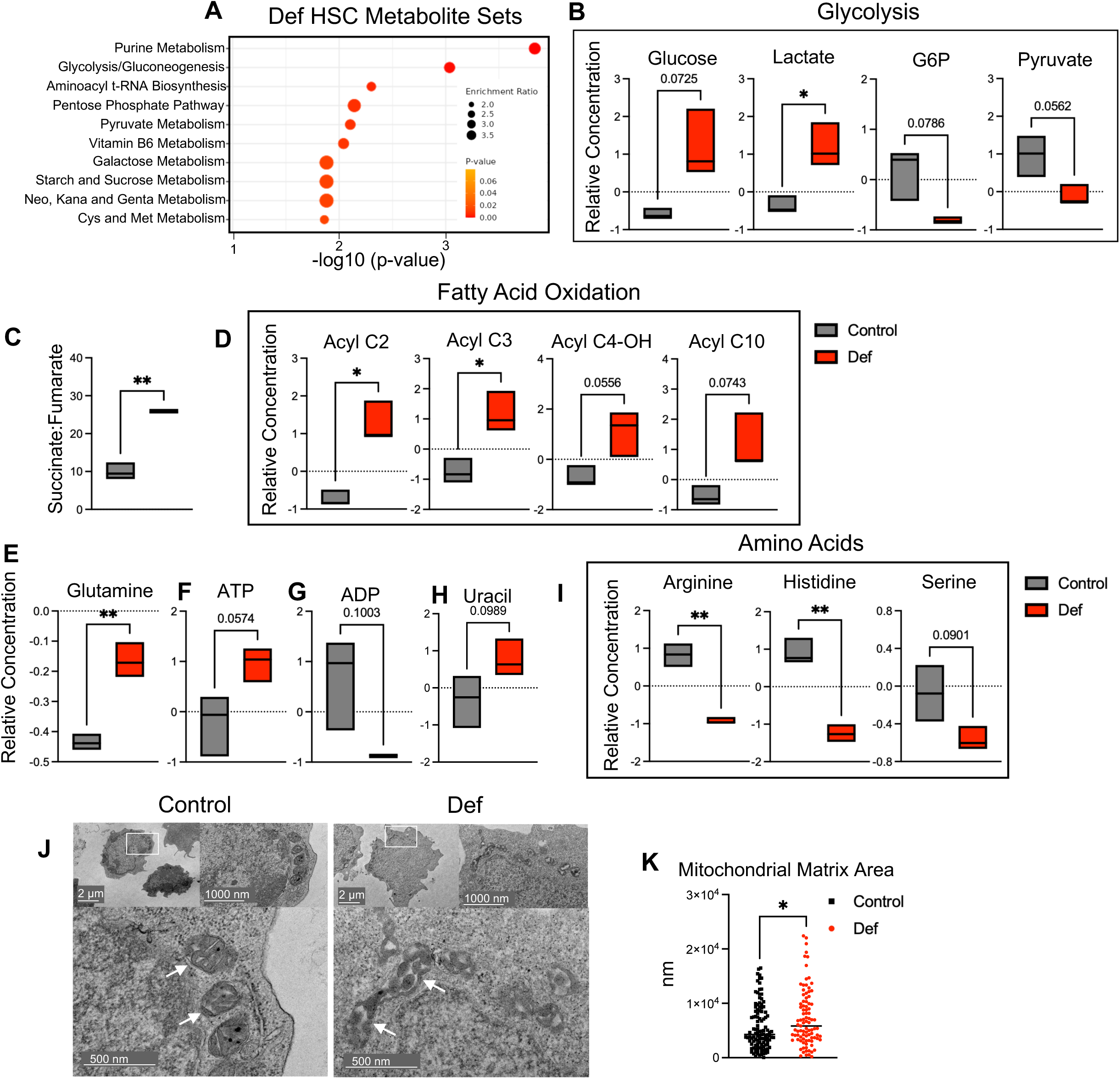
Maternal folate deficiency imprints a dysfunctional mitochondrial metabolic program in adult HSCs. **A)** Enriched metabolite sets in Def adult offspring HSCs determined by mass-spectrometry based metabolomic profiling. **B)** Normalized relative abundances of select individual metabolites within the glycolysis pathway identified by untargeted metabolomics analysis: Glucose, Lactate, Glucose-6-Phosphate, Pyruvate. **C)** Succinate:fumarate ratio based on relative abundances from untargeted metabolomics analysis. **D)** Normalized relative abundances of selected individual metabolites with the fatty acid oxidation pathways identified by untargeted metabolomics analysis: Acyl C2, Acyl C3, Acyl C4-OH, Acyl C10. **E)** Normalized relative abundance of glutamine identified by untargeted metabolomics. **F)** Normalized relative abundance of ATP identified by untargeted metabolomics. **G)** Normalized relative abundance of ADP identified by untargeted metabolomics. **H)** Normalized relative abundance of uracil identified by untargeted metabolomics. **I)** Normalized relative abundances of select individual metabolites within the aminoacyl t-RNA. biosynthesis pathway identified by untargeted metabolomics analysis: Arginine, Histidine, Serine. **J)** Representative electron microscopy images of Control and Def adult offspring HSCs. White box highlights region of interest. Images taken at 15,000x magnification. **K)** Inner mitochondrial space in Control and Def electron microscopy mitochondria measured by ImageJ, normalized by pixel. N = 3 independent experiments representing three mice from independent litters per group. Data represented as range of data points. Significance determined unpaired student’s t-test; *P < 0.05; **P < 0.01.

### Single-cell transcriptional profiling links folate deficient offspring metabolic phenotypes to stress programs

We performed single-cell RNA sequencing (scRNA-seq) to define transcriptional programs underlying metabolic dysfunction in folate Def offspring HSCs (**Fig. 3**). Unsupervised clustering identified LT-HSC, multipotent progenitor, and lineage-primed progenitor populations based on established markers (**Fig. 3A; Fig. S4A-B**). The greatest number of differentially expressed (DE) genes occurred within HSC clusters (**Fig. 3B**), consistent with early programming being driven at the HSC level. Def dormant (d)HSCs exhibited repression of mitochondrial programs, including oxidative phosphorylation (OXPHOS), electron transport chain (ETC) function, ATP synthesis, mitochondrial complex assembly and translation-related transcriptional programs (**Fig. 3C-D**). Expression of multiple ETC components spanning Complex I–IV were reduced (**Fig. 3E**). GSEA further demonstrated reduced OXPHOS, glycolysis, and fatty acid oxidation pathways in Def HSCs (**Fig. 3F**).

**Figure 3.**
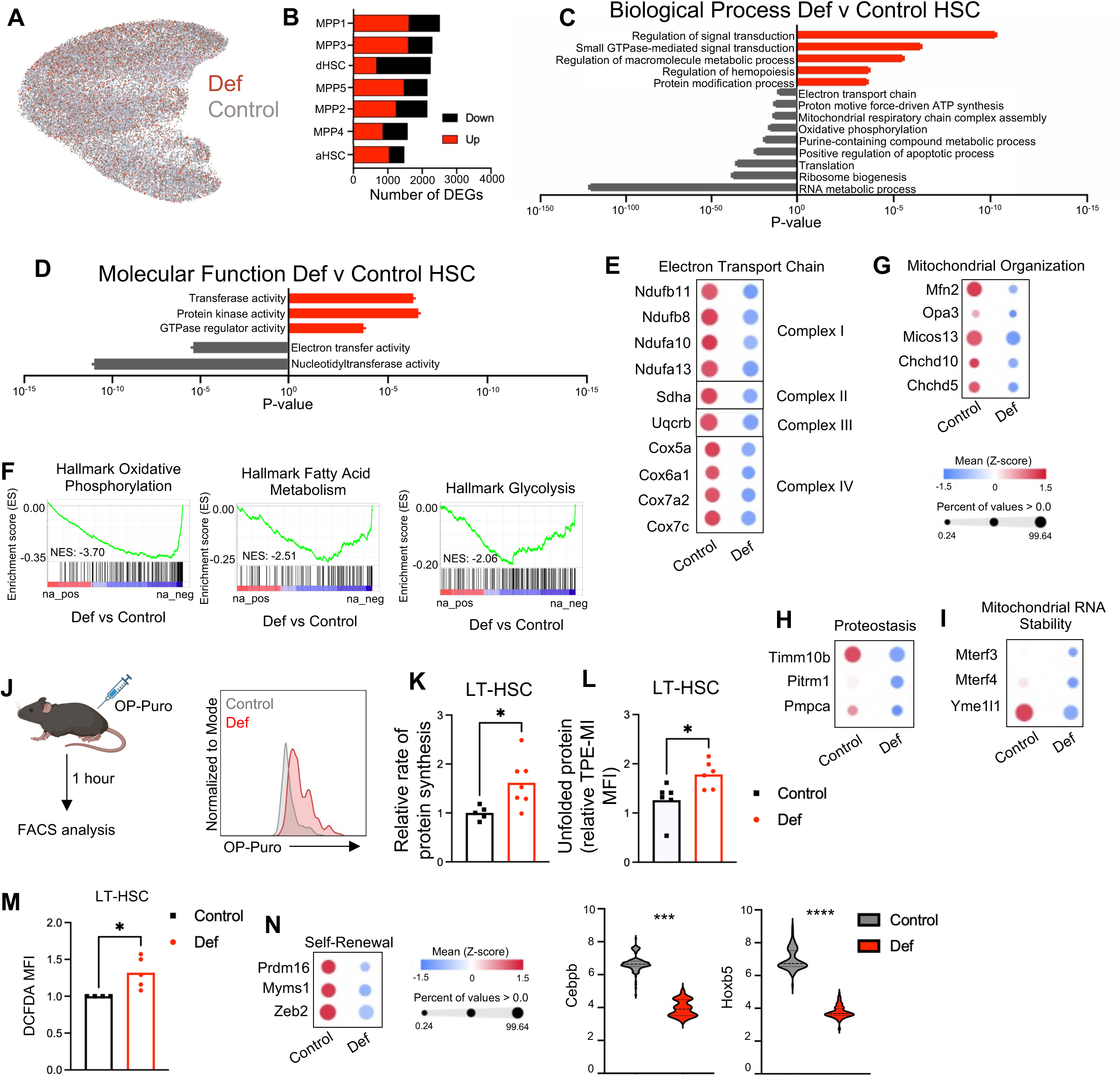
Single-cell transcriptional profiling links folate deficient adult offspring metabolic phenotypes to stress programs. **A)** UMAP projection representation of Def and Control populations identified by single-cell sequencing (scRNAseq). **B)** Number of differentially expressed genes per cluster in Def offspring HSCs, separated by down-regulated (black) and up-regulated (red) genes. **C)** Gene ontology biological process pathways in Def vs Control HSCs, separated by down-regulated (grey) and up-regulated (red) pathways. **D)** Gene ontology molecular functions pathways in Def vs Control HSCs, separated by down-regulated (grey) and up-regulated (red) pathways. **E)** Bubble plot representation of gene expression for electron transport genes in Complexes I-IV in Def and Control HSCs. **F)** Gene set enrichment analysis of downregulated hallmark metabolic pathways in Def HSCs. **G)** Bubble plot representation of gene expression for mitochondrial organization genes in Def and Control HSCs. **H)** Bubble plot representation of gene expression for proteostasis in Def and Control HSCs. **I)** Bubble plot representation of gene expression for mitochondrial RNA stability in Def and Control HSCs. **J)** OP-Puro experimental schematic, and representative histogram of average OP-Puro intensity. **K)** Relative rate of protein synthesis in Def and Control offspring LT-HSCs, measured by OP-Puro incorporation, normalized to internal wild-type control. **L)** Unfolded protein rates in Def and Control offspring LT-HSCs by tetraphenylethene maleimide (TPE-MI) mean fluorescence intensity. **M)** Measurement of intracellular reactive oxygen species in adult offspring Def and Control LT- and ST-HSCs by DCFDA ((2’,7’)-dichlorofluorescin diacetate) using flow cytometry. **N)** Bubble plot representation of gene expression for self-renewal genes in Def and Control HSCs alongside violin plots of Cebpb and Hoxb5 expression. N = 3 independent experiments representing three mice from independent litters per group. Data represented as range of data points. Significance determined by student’s t-test; *P < 0.05; ***P < 0.001; ****P < 0.0001

In addition to mitochondrial metabolic defects, Def HSCs exhibited reduced expression of genes involved in mitochondrial architecture and cristae organization, including Mfn2 and Opa3 ^62,63^ (**Fig. 3G**). Mfn2 coordinates multiple aspects of mitochondrial protein handling,^64^ and genes regulating mitochondrial protein import and proteostasis were also downregulated (**Fig. 3H**), along with mediators of mitochondrial RNA stability and inner membrane quality control^65,66,67^ (**Fig. 3I**). To assess functional consequences of altered proteostasis, we measured in vivo protein synthesis using OP-Puro incorporation (**Fig. 3J**).^68^ Surprisingly, Def HSCs exhibited increased protein synthesis (**Fig. 3K**) but elevated unfolded protein burden (**Fig. 3L**). ^69^ We also observed elevated ROS levels in Def HSCs (**Fig. 3M**). These findings demonstrate that maternal folate deficiency drives a proteostatic imbalance in which translational output outpaces quality-control capacity.

Given the requirement for tightly regulated protein synthesis and mitochondrial fitness to maintain LT-HSC quiescence,^30,37^ metabolic dysregulation would be expected to compromise transcriptional programs governing self-renewal. Accordingly, core regulators of HSC identity and maintenance^70–73^ were downregulated in Def HSCs (**Fig. 3N**). These findings suggest that impaired mitochondrial homeostasis converges on regulatory networks governing self-renewal in Def offspring HSCs.

### Maternal folate supplementation enhances mitochondrial fitness and bioenergetic capacity in offspring HSCs

We performed metabolomic analysis to further investigate the enhanced self-renewal phenotype in Supp offspring HSCs. Pathway analysis of the five differentially abundant metabolites identified sulfur and purine metabolism as the primary altered pathways (**Fig. 4A**). Aspartate and GTP were reduced in Supp HSCs (**Fig. 4B-C**), although GTP was not reliably detected by untargeted LC-MS and therefore requires validation. Reduced aspartate levels were consistent with increased anaplerotic utilization,^74^ whereas reduced pyruvate levels suggested enhanced mitochondrial utilization (**Fig. 4D**).

**Figure 4.**
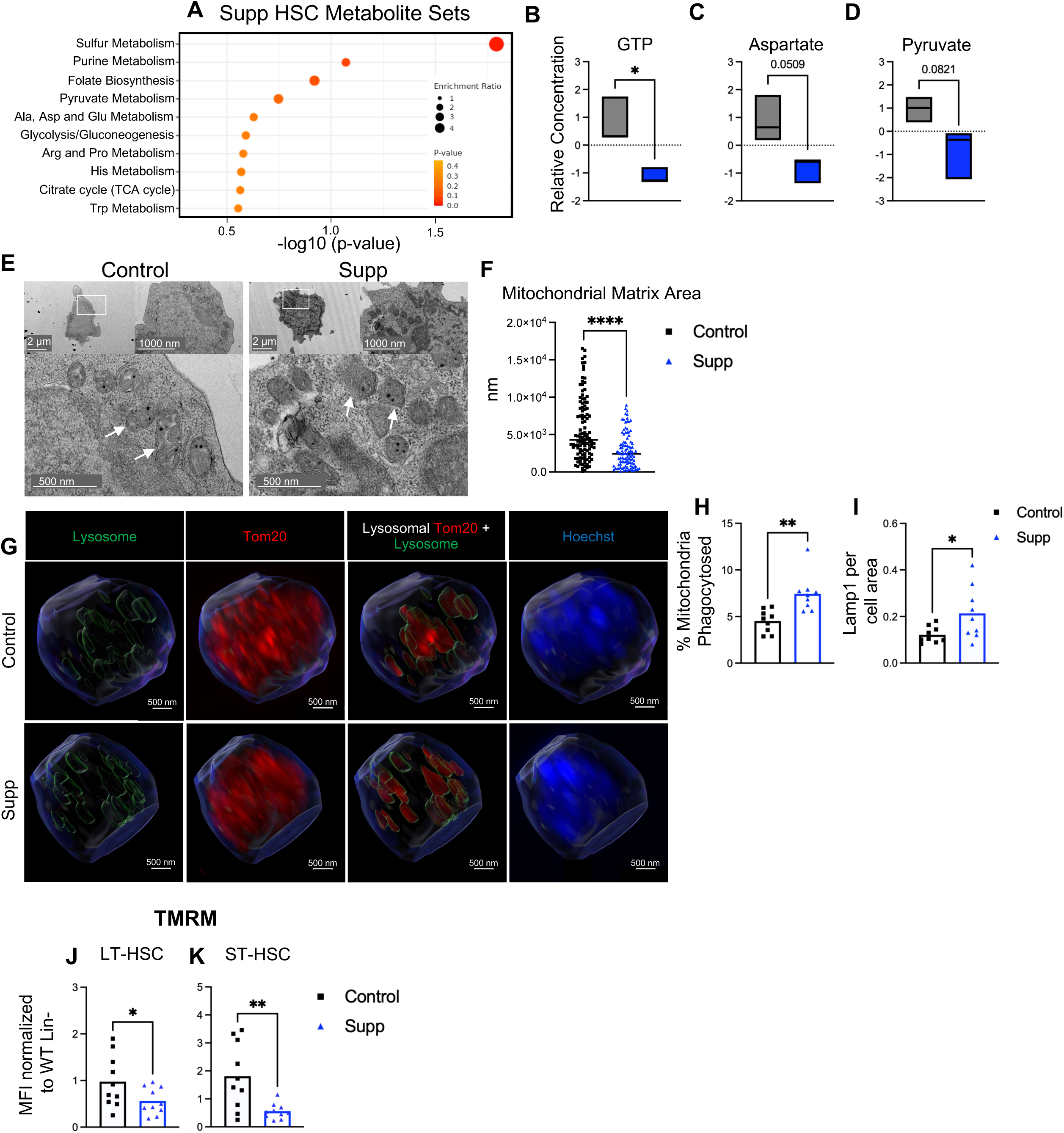
Maternal folate supplementation enhances mitochondrial fitness and bioenergetic capacity in adult HSCs. **A)** Enriched metabolite sets in Supp adult offspring HSCs determined by mass-spectrometry based metabolomic profiling. **B-D)** Normalized relative abundances of selected individual metabolites identified by untargeted metabolomics analysis: (**B**) GTP (**C**) Aspartate (**D**) Pyruvate. **E)** Representative electron microscopy images of Control and Supp adult offspring HSCs, white box indicates enhanced region. Images taken at 15,000x magnification. **F)** Inner mitochondrial space in Control and Supp electron microscopy mitochondria measured by ImageJ, normalized by pixel. **G)** Imaris 3D reconstruction of super-resolution confocal microscopy images on sorted Supp and Control adult offspring HSCs, stained for Tom20 (red), Lamp1 (green), DAPI (blue) and reconstructed lysosome organelles (green). Intracellular TOM20 staining was used to reconstruct the cell surface and intracellular mitochondrial compartment. LAMP1 staining was used to reconstruct lysosomes. **H)** Percent of mitochondria present inside reconstructed lysosomes as a percentage of total mitochondria in Supp and Control adult offspring HSCs. **I)** Total Lamp1 signal intensity as determined by super-resolution confocal microscopy by cell area (um^2^) in Supp and Control adult offspring HSCs. **J-K)** Mean fluorescence intensity (MFI) of tetramethylrhodamine (TMRM) in (**K**) LT-HSC and (**L**) ST-HSC Supp and Control adult offspring populations, normalized to internal wild-type control. N = 3 independent experiments representing three mice from independent litters per group. Data represented as range of data points. Significance determined by unpaired student’s t-test; *P < 0.05; **P < 0.01; ****P < 0.0001

Given the importance of GTPase-mediated control of mitochondrial dynamics,^75^ we examined mitochondrial morphology by TEM. Supp HSCs exhibited more densely packed cristae (**Fig. 4E; Fig. S3E**) and reduced inner mitochondrial space (**Fig. 4F**), associated with improved mitochondrial respiratory efficiency.^60,61^ We performed Airyscan microscopy on sorted LT-HSCs labeled for mitochondria and lysosomes (**Fig. S5A**) to examine mitochondrial morphology. Supp HSCs exhibited increased mitochondrial material within LAMP1⁺ lysosomal compartments (**Fig. 4G-H**). Importantly, this increase was not attributable to elevated mitochondrial content (**Fig. S5B**), indicating a true enhancement of mitochondrial turnover. Supp HSCs also exhibited increased LAMP1⁺ signal intensity (**Fig. 4I**), suggesting expansion of the lysosomal compartment. To determine whether these structural changes translated to altered mitochondrial function, we measured mitochondrial membrane potential using TMRM. Supp HSCs exhibited decreased TMRM intensity (**Fig. 4J-K**), indicating reduced mitochondrial activity. Together, these data provide direct imaging-based evidence that maternal folate supplementation promotes lysosome-dependent mitochondrial turnover, establishing a distinct mitochondrial phenotype in adult HSCs characterized by enhanced quality control and reduced mitochondrial activity rather than global metabolic rewiring.

### Single-cell transcriptional states are associated with enhanced mitochondrial fitness and self-renewal in folate supplemented offspring HSCs

To determine whether enhanced mitochondrial fitness in folate Supp offspring HSCs reflects coordinated transcriptional programs, we performed scRNA-seq (**Fig. 5A**). Transcriptional programming was most pronounced in HSC subsets (**Fig. 5B**). Pathway enrichment in dHSCs revealed upregulation of G protein signaling, autophagy, autophagosome formation, and cellular component organization (**Fig. 5C).** Molecular function analysis further identified GTP binding and GTPase activity among the dominant upregulated categories (**Fig. 5D)**. Supp HSCs upregulated small GTPases and their regulators (**Fig. 5E-F**), including Rab-family GTPases, GTPase regulators, and signaling mediators such as Kras.^76–78^ Supp HSCs upregulated genes supporting mitochondrial architecture (**Fig. 5G**), as well as genes associated with autophagy and mitophagy (**Fig. 5H**), suggesting reinforcement of cristae integrity^79–81^ and enhanced organelle surveillance.^82–87^ Asxl2, Ikzf2, and other regulators of long-term self-renewal and chromatin integrity^88–90^, were also upregulated (**Fig. 5I**). Collectively, these changes suggest that GTPase signaling links enhanced autophagy with improved mitochondrial quality control to support a coordinated program of self-renewal.

**Figure 5.**
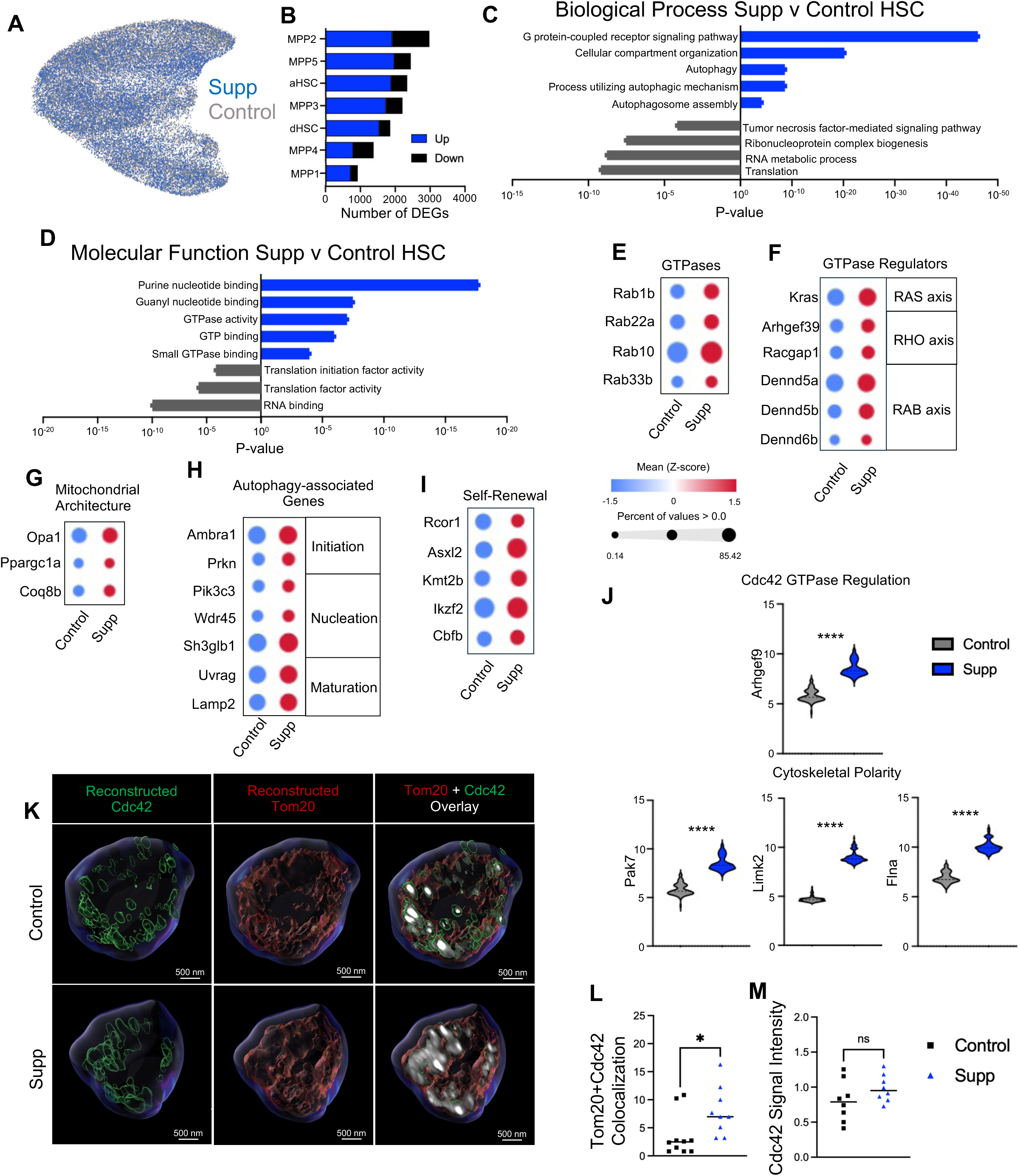
Single-cell transcriptional states are associated with enhanced mitochondrial fitness and self-renewal in folate supplemented adult offspring HSCs. **A)** UMAP projection representation of Supp and Control populations identified by single-cell sequencing (scRNAseq). **B)** Number of differentially expressed genes per cluster in Supp offspring HSC seSeq, separated by down-regulated (black) and up-regulated (blue) genes. **C)** Gene ontology biological process pathways in Supp vs Control HSCs, separated by down-regulated (grey) and up-regulated (blue) pathways. **D)** Gene ontology molecular functions pathways in Supp vs Control HSCs, separated by down-regulated (grey) and up-regulated (blue) pathways. **E)** Bubble plot representation of gene expression for GTPase genes in Supp and Control offspring HSCs. **F)** Bubble plot representation of gene expression for GTPase regulator genes in Supp and Control offspring HSCs. **G)** Bubble plot representation of gene expression for mitochondrial architecture genes in Supp and Control offspring HSCs. **H)** Bubble plot representation of gene expression for autophagy-associated genes in Supp and Control offspring HSCs. **I)** Bubble plot representation of gene expression for self-renewal genes in Supp and Control offspring HSCs. **J)** Violin plot representation of CDC42-interacting genes in Control and Supp offspring HSCs, based on GTP regulation or cytoskeletal polarity. **K)** Imaris 3D reconstruction of super-resolution confocal microscopy images on sorted Supp and Control offspring HSCs, stained and reconstructed for Tom20 (red), CDC42-GTP (green), Hoechst (blue). 3D Reconstruction includes Tom20+CDC42-GTP overlap (white). **L)** Colocalization of Tom20 (red) mitochondrial 3D reconstruction with CDC42-GTP (green) 3D reconstruction in Supp and Control offspring HSCs within regions of interest, represented as percentage. **M)** CDC42-GTP signal intensity by super-resolution confocal microscopy by cell volume in Supp and Control offspring HSCs (μm^3^). N = 3 independent experiments representing three mice from independent litters per group. Data represented as range of data points. Significance determined by unpaired student’s t-test; *P < 0.05; ****P < 0.0001

Small GTPases, particularly CDC42, are central determinants of HSC polarity ^91^ and upregulation of CDC42-interacting genes including Arhgef9,^92^ Pak7,^93^ Limk2,^94^ and Flna^95^ (**Fig. 5J**) suggests that enhanced GTPase signaling alters subcellular organelle positioning to support improved mitochondrial quality control in Supp HSCs. We performed Airyscan imaging of CDC42 and mitochondria in sorted HSCs (**Fig. S5C**) and identified increased mitochondrial-associated CDC42 signal in Supp HSCs (**Fig. 5K-L**). Total CDC42 signal intensity was unchanged between Supp and Control HSCs (**Fig. 5M**), indicating this enrichment reflects spatial redistribution rather than increased protein abundance. Overall, Supp HSCs exhibited enhanced CDC42-associated spatial clustering of mitochondria, consistent with enhanced organelle quality control.

### Maternal folate availability establishes distinct metabolic programs in fetal HSCs

Because adult HSC phenotypes persist despite normalization of postnatal folate intake (**Fig. S1O**), we hypothesized that maternal folate availability alters the developing fetal hematopoietic landscape. Maternal folate deficiency increased total FL CD45⁺ cells, whereas supplementation reduced CD45⁺ cells (**Fig. 6A**). Def fetuses exhibited expansion of LT- and ST-HSCs and lymphoid-biased drHSCs (**Fig. 6B-D**), similar to prenatal inflammation.^46^ MPP2^MkE^ increased in Def fetuses (**Fig. S6A**), while other MPPs were unchanged (**Fig. S6B-C**). CD11b⁺ cells increased in Def and decreased in Supp FLs (**Fig. S6D**), with no changes in FL B cells (**Fig. S6E**). Seahorse analysis revealed reduced basal glycolytic capacity in folate Def fetal HSPCs (**Fig. 6E-F; Fig. S6F**), whereas Supp fetal HSPCs exhibited increased glycolytic capacity (**Fig. 6E-F; Fig. S6F**) and elevated basal and maximal oxygen consumption rates (**Fig. 6G-H; Fig. S6G**). TMRM staining further demonstrated increased mitochondrial membrane potential (MMP) in Def fetal HSCs and MPPs, and to a lesser extent in Supp fetal HSCs (**Fig. S6H-J**), indicating folate-dependent modulation of fetal HSPC metabolic state.

**Figure 6.**
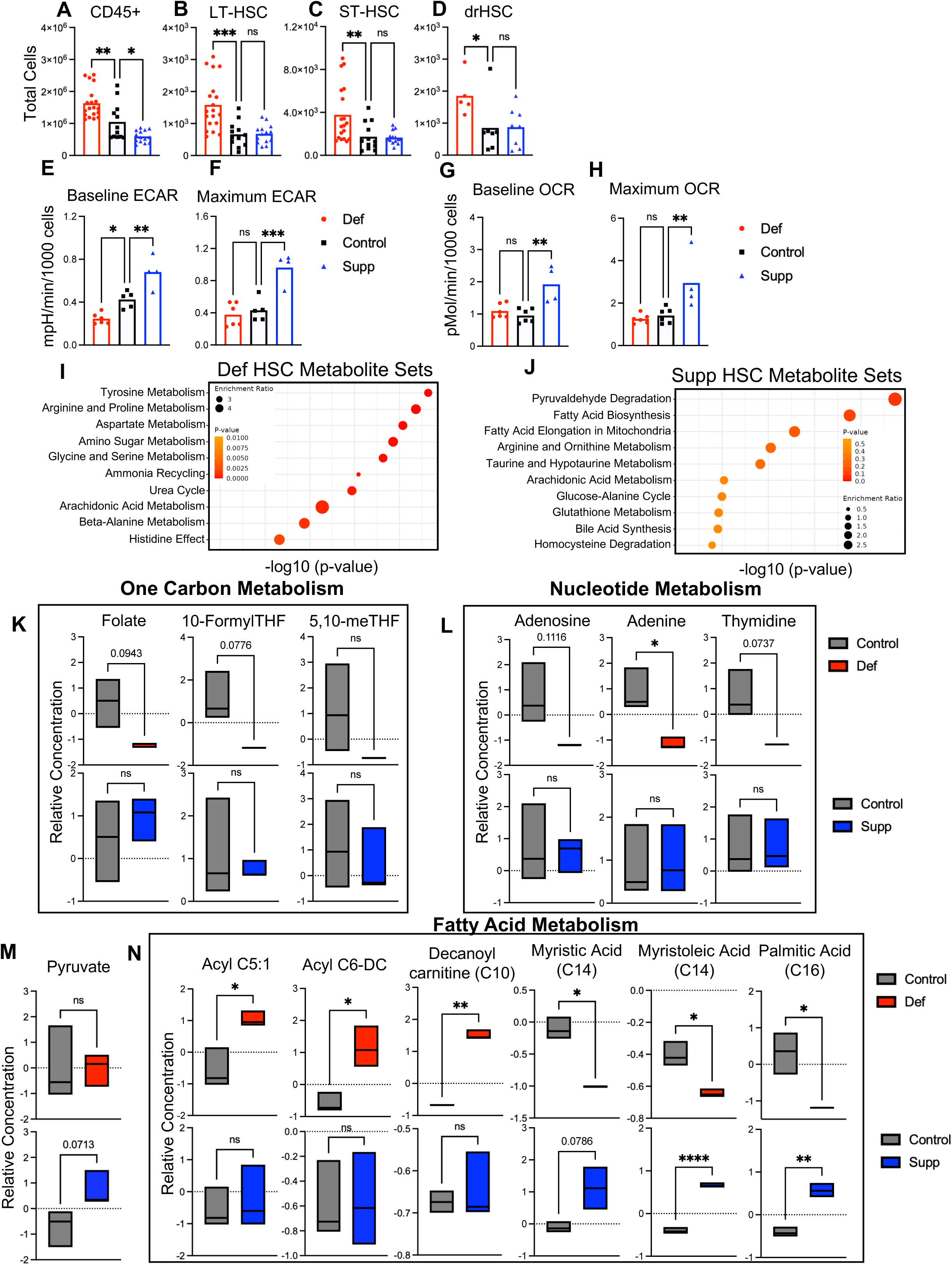
Maternal folate availability establishes distinct metabolic programs in fetal HSCs. **A-D)** Total cellularity of fetal liver hematopoietic cell populations quantified by flow cytometry. (**A**) CD45^+^ cells, (**B**) LT-HSC, (**C**) ST-HSC, (**D**) developmentally-restricted HSC (drHSC). **E-F**) Fetal liver HSPC baseline, (**E**) and maximal (**F**) extracellular acidification rate by Seahorse metabolic assay, normalized to 1000 cells per well. **G-H**) Fetal liver HSPC baseline, (**G**) and maximal (**H**) oxygen consumption rate by Seahorse metabolic assay, normalized to 1000 cells per well. **I)** Enriched metabolite sets in Def fetal liver HSCs determined by mass-spectrometry based metabolomic profiling. **J)** Enriched metabolite sets in Supp fetal liver HSCs determined by mass-spectrometry based metabolomic profiling. **K)** Normalized relative abundances of select individual metabolites in Def, Control and Supp fetal liver HSCs within the one-carbon metabolism pathway identified by untargeted metabolomics analysis: Folate, 10-FormylTHF, 5,10-methylTHF. **L)** Normalized relative abundances of select individual metabolites in Def, Control and Supp fetal liver HSCs within the nucleotide metabolism pathway identified by untargeted metabolomics analysis: Adenosine, Adenine, Thymidine. **M)** Normalized relative abundance of pyruvate identified by untargeted metabolomics analysis. **N)** Normalized relative abundances of select individual metabolites in Def, Control and Supp fetal liver HSCs within the fatty acid metabolism pathway identified by untargeted metabolomics analysis: Acyl C5:1, Acyl C6-DC, decanoyl carnitine (C10), myristic acid (C14), myristoleic acid (C14) and palmitic acid (C16). N = 3 independent experiments representing three mice from independent litters per group. Data represented as range of data points. Significance determined by one-way ANOVA with multiple comparisons or unpaired student’s t-test for analyses with two treatment groups; *P < 0.05; ** P < 0.01; ***P < 0.001; ****P < 0.0001

To further contextualize these bioenergetic phenotypes, we examined fetal HSC metabolism by unbiased metabolomics at E14.5. Folate Def fetal HSCs showed the most distinct metabolic profile compared with Control and Supp groups (**Fig. S7A; Fig. 6I-J**), with particularly pronounced changes in amino acid metabolism (**Fig. 6I**). Def fetal HSCs exhibited reduced one-carbon and nucleotide metabolites (**Fig. 6K-L**), alongside reduced glucose and increased glutamine and glutamate (**Fig. S7B**). Methionine S-oxide accumulation relative to methionine further indicated increased oxidative pressure (**Fig. S7C**). In contrast, pyruvate was selectively increased in Supp fetal HSCs (**Fig. 6M**).

Maternal folate drove opposing alterations to lipid metabolism, with Def fetal HSCs exhibiting increased short- and odd-chain acylcarnitines (**Fig. 6N**) and reduced long-chain fatty acids (**Fig. 6N; Fig. S7D**) consistent with altered fatty acid processing and incomplete mitochondrial β-oxidation.^57,58^ In contrast, Supp fetal HSCs showed increased long-chain fatty acids and enrichment of fatty acid biosynthesis pathways (**Fig. 6J, N**), suggesting enhanced lipid availability rather than global lipid remodeling. Collectively, these data reveal that maternal folate status induces metabolic stress in Def fetal HSCs and more selective metabolic tuning in Supp fetal HSCs that may support mitochondrial function without inducing metabolic stress.

### Prenatal folate nutrition initiates long-term remodeling of chromatin accessibility in adult hematopoietic stem cells

To assess whether maternal folate availability leaves a lasting imprint on chromatin accessibility, we performed ATAC-seq on adult offspring HSCs (**Fig. 7**). Principal component analysis separated all three groups (**Fig. S8A**). Differential accessibility analysis showed that folate deficiency was associated with open chromatin regions (**Fig. 7A; Fig. S8B**), whereas supplementation was predominantly associated with closed regions (**Fig. 7B; Fig. S8C**). Supp HSCs showed promoter enrichment in open chromatin, whereas Def HSCs showed promoter enrichment in closed chromatin (**Fig. S8D-E**). Pathway enrichment of differentially accessible regions (DARs) identified programs consistent with single-cell transcriptional profiling (**Fig. 7C-D**), supporting a relationship between altered chromatin accessibility and sustained transcriptional programming.

**Figure 7.**
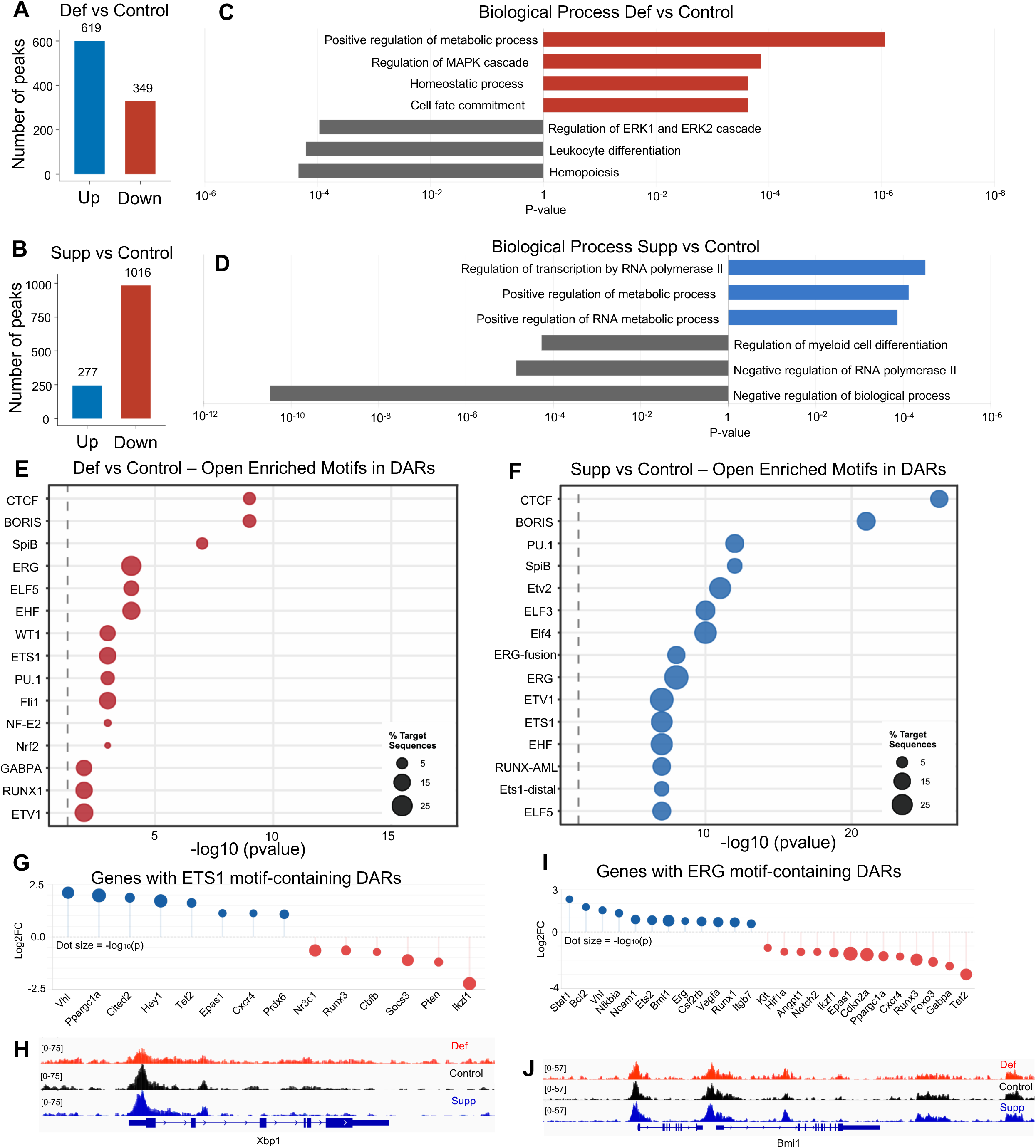
Prenatal folate nutrition drives long-term remodeling of chromatin accessibility in adult HSCs. **A-B)** Number of open and closed differentially accessible regions (DARs) in Def vs Control (**A**) and Supp vs Control (**B**) adult offspring HSCs. **C-D)** Enriched gene ontology pathways in open and closed DARs in Def vs Control (**C**) and Supp vs Control (**D**) adult offspring HSCs. **E-F)** Enriched transcription factor (TF) binding motifs in open DARs in Def vs Control (**E**) and Supp vs Control (**F**) adult offspring HSCs. **G)** Genes associated with ETS1 motif-containing DARs in Def vs Control HSCs. **H)** IGV gene track of Xbp1 DAR within promoter in adult offspring HSCs. **I)** Genes with ERG motif-containing DARs in Supp vs Control adult offspring HSCs. **J)** IGV gene track of Bmi1 DAR within distal enhancer in adult offspring HSCs. N = 3 independent experiments representing three mice from independent litters per group.

Motif enrichment analysis identified differentially accessible transcription factor (TF) binding sites in Def and Supp HSCs. Def HSCs showed increased accessibility of PU.1 and ETS1 motifs (**Fig. 7E**), known regulators of hematopoietic lineage specification ^96^ and stem cell maintenance, respectively. Consistent with ETS1-associated enrichment, Cited2, a downstream target,^97^ and Epas1 (encoding Hif2α), a known ETS1-interacting gene,^98^ showed increased chromatin accessibility (**Fig. 7G**). Nrf2 and NF-E2 motifs were also enriched, with increased accessibility at Prdx6 (**Fig. 7E; Fig. S8H**), a target of Nrf2-mediated antioxidant regulation.^99^ Xbp1, a critical regulator of the unfolded protein response, oxidative stress adaptation^100^, and self-renewal^101^ showed decreased promoter accessibility (**Fig. 7H**). Collectively, these data suggest that chromatin remodeling in Def HSCs reflects metabolic rewiring in response to cellular stress.

TF motif enrichment of Supp HSCs identified increased accessibility of ERG- and ETS-family TF motifs (**Fig. 7F**). Increased accessibility at downstream target loci of ERG (**Fig. 7I**), a master regulator of HSC maintenance and self-renewal,^102,103^ included a distal enhancer regulating the Polycomb repressor Bmi1 (**Fig. 7I-J**), with reduced accessibility at its target genes Cdkn2a and Cdkn2b ^104^ (**Fig. 7I; Fig. S8I**). Polycomb-associated components Sfmbt1 ^105^ and Rybp ^106^ also showed increased accessibility (**Fig. S8I**). Loci within other self-renewal genes, Sall4 and Bcl2, ^107,108^ were also more accessible (**Fig. S8J**). Among metabolic genes, Pdk4, a regulator of mitochondrial pyruvate import,^109^ and Hk2, which catalyzes the first committed step of glycolysis,^110^ showed increased accessibility (**Fig. S8K**), suggesting coordinated chromatin remodeling of glucose and pyruvate handling. Collectively, maternal folate supplementation establishes a chromatin landscape favoring self-renewal and maintenance.

CTCF target loci were among the most significantly enriched motifs in both open and closed DARs across Def and Supp HSCs (**Fig. 7E-F; Fig. S8F-G**). Notably, the direction of enrichment was condition-specific: CTCF motifs were enriched in open regions in Supp HSCs (**Fig. 7F**), whereas CTCF was the top-ranked motif among closed regions in Def HSCs (**Fig. S8F**). Given the central role of CTCF in chromatin looping,^111^ insulation,^112^ and higher-order genome organization,^113,114^ as well as HSC self-renewal,^115^ these condition-specific patterns suggest that maternal folate availability influences accessibility at genomic regions associated with CTCF-mediated chromatin regulation. Overall, maternal folate availability remodels chromatin accessibility landscapes in offspring HSCs through divergent regulatory pathways that may shape lifelong hematopoietic function.

## DISCUSSION

Our findings demonstrate that maternal manipulation of a single micronutrient, folic acid, during development is sufficient to program adult HSC function across metabolic, transcriptional, and chromatin regulatory layers. This work extends prior research showing that environmental perturbations during development can alter offspring HSC function ^23,24,46,116^ by demonstrating that maternal folate availability during fetal hematopoiesis induced lasting alterations to both stem cell metabolism and function. Unlike adult HSCs, where metabolic perturbations are often reversible or context-dependent,^32,34,117^ fetal HSCs are actively cycling and metabolically distinct from their quiescent adult counterparts,^118^ raising the possibility that transient metabolic inputs may be encoded into persistent stem cell programs. Our findings demonstrate that developing HSCs are uniquely sensitive to metabolic inputs during development.

A recurring theme was the divergence of mitochondrial phenotypes in adult HSCs following maternal folate deficiency versus supplementation. This suggests that mitochondrial programming is not simply a function of deviation from an optimal folate set point but is differentially shaped by the metabolic consequences of folate restriction versus excess. Folate-dependent OCM supports nucleotide biosynthesis, cellular methylation, redox balance and resistance to oxidative stress.^19,21,119^ Thus, deviations in folate status may engage distinct compensatory programs, positioning mitochondrial homeostasis as a central, bidirectionally sensitive node through which developmental folate exposure shapes HSC function.

Several lines of evidence point toward altered mitochondrial quality control as a shared underlying axis of divergent mitochondrial programming. Def HSCs displayed broad suppression of oxidative phosphorylation and disrupted cristae, consistent with impaired mitochondrial homeostasis. Def HSCs also suppressed Mfn2, which maintains mitochondrial network integrity, and modulates mitochondrial ultrastructure.^120^ Mfn2 is required for HSC self-renewal and lymphoid potential,^121^ consistent with functional deficits observed in Def HSCs. Conversely, Supp HSCs exhibited greater lysosomal engulfment of mitochondria, accompanied by transcriptional enrichment of autophagy-associated pathways, suggesting that enhanced respiratory efficiency and efficient organelle turnover contributed to stem cell fitness. Given evidence implicating CDC42 in organelle polarity and lysosome-mediated organelle quality control,^122,123^ increased mitochondrial localization of CDC42 in Supp HSCs may contribute mechanistically to the enhanced mitochondrial quality-control phenotype by maternal folate supplementation.

Modulation of maternal folate nutrition elicited inverse effects on fetal liver cellularity, likely reflecting contrasting metabolic responses rather than a simple dose-response relationship. We propose that stress-mediated expansion of fetal HSCs under folate deficiency compromises long-term self-renewal, whereas the more restrained response to supplementation preserves a higher-quality stem cell pool. A central question raised by our findings is how transient fetal metabolic states become encoded into durable adult HSC programs. One candidate mechanism is direct coupling between metabolite availability and chromatin regulation, as methylation and acetylation depend on metabolite intermediates such as S-adenosylmethionine (SAM) and acetyl-CoA.^124–126^ Although SAM was not reliably detected, folate Def fetal HSCs showed reduced one-carbon intermediates and increased methionine S-oxide, indicative of impaired methylation potential and elevated oxidative pressure.^127^ These findings provide a plausible substrate-level link between fetal metabolic insufficiency and suppressed chromatin accessibility at mitochondrial and stress-response loci observed in adult Def HSCs. In Supp HSCs, our data implicate pyruvate availability as a potential substrate linking metabolic state to chromatin regulation and membrane-associated signaling.^128^ Pyruvate-derived acetyl-CoA supports histone acetylation, and nuclear pyruvate dehydrogenase complexes generate acetyl-CoA for epigenetic regulation.^129^ Although we could not directly detect acetyl-CoA, increased fetal pyruvate is consistent with greater substrate availability for acetylation-dependent chromatin regulation. Increased Pdk4 accessibility in adult Supp HSCs, a regulator of mitochondrial pyruvate flux,^109^ raises the possibility that fetal pyruvate availability and adult chromatin accessibility are linked components of the same folate-dependent programming axis.

HSCs are known to be particularly sensitive to oxidative and proteotoxic stress and rely on tightly restricted rates of protein synthesis to protect long-term function.^37,130^ In Def HSCs, motif enrichment analysis identified accessible regulatory elements associated with metabolic stress responses, alongside reduced accessibility at the Xbp1 promoter. Together, these signatures suggest that maternal folate deficiency establishes a stress-adapted epigenetic state favoring short-term metabolic compensation at the expense of homeostatic maintenance programs required for self-renewal. These chromatin signatures were also reflected in impaired proteostasis and increased oxidative stress observed in Def adult HSCs. Given the role of Polycomb proteins as an epigenetic checkpoint coupling cell-cycle control to HSC self-renewal,^131,132^ increased accessibility at a Bmi1 enhancer in Supp HSCs is notable, and coincided with increased accessibility at stem cell maintenance programs identified by scRNA-seq. Thus, developmental metabolic perturbations can encode persistent chromatin-level changes that alter HSC metabolism and function. Notably, the divergence in enrichment of CTCF motifs by maternal folate status may suggest remodeling of higher order genome architecture to regulate long term HSC function and metabolism.

Together, these findings reveal that maternal folate availability is metabolically written into developing HSCs and locked in at the chromatin level, converting a transient developmental state into heritable cellular memory. While metabolic regulation of stem cell fate has been studied primarily in adult HSCs, ^29,133–135^ our findings extend this paradigm to fetal development and highlight maternal nutrition as a determinant of HSC function. While common polymorphisms in folate-mediated OCM genes yield functional folate deficiency despite adequate intake and mandatory fortification,^7–10^ our findings raise the possibility that genetic susceptibility could drive similar HSC programming irrespective of maternal folate status. Whereas maternal folate deficiency impaired long-term HSC function, enhanced self-renewal following supplementation should not be interpreted as unequivocally beneficial. Circulating unmetabolized folic acid associated with excess folate intake^11–13^ is causally associated with cancer prevalence, ^13,136–140^ though mechanisms remain unknown. Enhanced self-renewal may therefore carry risks, particularly in the context of age- or mutation-associated clonal expansion. As vitamin supplement use rises ^141,142^ alongside mandatory folic acid fortification, the long-term consequences of excess micronutrient supplementation for aging HSCs may not be evident for another 10-20 years. Our findings identify maternal nutrient availability as an underappreciated determinant of adult HSC metabolism and function, with implications for prenatal nutrition and population-wide folate fortification.

## Supporting information

All Supplemental Materials

## Acknowledgements

The authors would like to thank James Marvin and the University of Utah Flow Cytometry Core, and the University of Utah Comparative Medicine Center. Research reported in this publication utilized the Health Sciences Cores Cell Imaging Core at the University of Utah and the High-Throughput Genomics and Cancer Bioinformatics Shared Resource at Huntsman Cancer Institute supported by the National Cancer Institute Award P30CA042014. Figures were generated using Biorender. We thank Drs. Diego López, Patrick Stover, and Luis Batista for critical reading of our manuscript. This work was supported by an NIH/National Heart Lung Blood Institute (NHLBI) Award R01HL157794 to A.E.B. and the Pew Biomedical Scholars award to A.E.B. A.E.B. is also the recipient of an NHLBI K01HL130753. B.P.K. is supported by Heart Lung Blood Institute (NHLBI) Award Disorders (NIDCD) training award F31 HL170732. L.F.C. was supported by the NIH National Institute of Diabetes and Digestive and Kidney Diseases (NIDDK) under award T32-DK007158. T.N. was supported by the NIH NHLBI award R01HL146442. F.J.Z was supported by NIH/NHLBI (F31HL170531) and NIH/NCI (2T32CA067754). R.A.J.S is supported by NIH/NIA (R01AG088725), NIH/NHLBI (R01HL180754), and the UC San Diego Sanford Stem Cell Institute Discovery Center. R.S.W., A.M.F, and B.M.C. were supported by the Mark Foundation Endeavor Award and A.M.F. is supported by the UAB T32 Training Program, T32AR069516.

## Author contributions

AEB and BPK designed the study; AEB, BPK, AN, RAJS, MSF, AC, TN, AD, RSW, and RL developed the methodology; BPK, AN, VC, BC, NH, FZ, LFC, AF, SA, and TN performed experiments; AEB, BPK, BC, RW, AN, AC, and TN analyzed and interpreted data; AEB obtained funding and supervised the study; AEB managed the project; AEB and BPK wrote the manuscript; and AEB, BPK, RSW, RAJS, TN, and MSF reviewed and edited the manuscript.

## REFERENCES

1. Prevention of neural tube defects: results of the Medical Research Council Vitamin Study. MRC Vitamin Study Research Group. (1991). Lancet 338, 131–137.

2. Use of folic acid for prevention of spina bifida and other neural tube defects--1983-1991. (1991). MMWR Morb Mortal Wkly Rep 40, 513–516.

3. Folic acid for the prevention of neural tube defects: U.S. Preventive Services Task Force recommendation statement. (2009). Ann Intern Med 150, 626–631. 10.7326/0003-4819-150-9-200905050-00009.

4. Recommendations for the use of folic acid to reduce the number of cases of spina bifida and other neural tube defects. (1992). MMWR Recomm Rep 41, 1–7.

5. Rogers, L.M., Cordero, A.M., Pfeiffer, C.M., Hausman, D.B., Tsang, B.L., De-Regil, L.M., Rosenthal, J., Razzaghi, H., Wong, E.C., Weakland, A.P., and Bailey, L.B. (2018). Global folate status in women of reproductive age: a systematic review with emphasis on methodological issues. Ann N Y Acad Sci 1431, 35–57. 10.1111/nyas.13963.

6. Pfeiffer, C.M., Sternberg, M.R., Zhang, M., Fazili, Z., Storandt, R.J., Crider, K.S., Yamini, S., Gahche, J.J., Juan, W., Wang, C.Y., et al. (2019). Folate status in the US population 20 y after the introduction of folic acid fortification. Am J Clin Nutr 110, 1088–1097. 10.1093/ajcn/nqz184.

7. Pereira, A.C., Schettert, I.T., Morandini Filho, A.A., Guerra-Shinohara, E.M., and Krieger, J.E. (2004). Methylenetetrahydrofolate reductase (MTHFR) c677t gene variant modulates the homocysteine folate correlation in a mild folate-deficient population. Clin Chim Acta 340, 99–105. 10.1016/j.cccn.2003.09.016.

8. Vidmar Golja, M., Šmid, A., Karas Kuželički, N., Trontelj, J., Geršak, K., and Mlinarič-Raščan, I. (2020). Folate Insufficiency Due to MTHFR Deficiency Is Bypassed by 5-Methyltetrahydrofolate. J Clin Med 9. 10.3390/jcm9092836.

9. Coppedè, F., Stoccoro, A., Tannorella, P., Gallo, R., Nicolì, V., and Migliore, L. (2019). Association of Polymorphisms in Genes Involved in One-Carbon Metabolism with MTHFR Methylation Levels. Int J Mol Sci 20. 10.3390/ijms20153754.

10. Li, W.X., Cheng, F., Zhang, A.J., Dai, S.X., Li, G.H., Lv, W.W., Zhou, T., Zhang, Q., Zhang, H., Zhang, T., et al. (2017). Folate Deficiency and Gene Polymorphisms of MTHFR, MTR and MTRR Elevate the Hyperhomocysteinemia Risk. Clin Lab 63, 523–533. 10.7754/Clin.Lab.2016.160917.

11. Obeid, R., Kirsch, S.H., Dilmann, S., Klein, C., Eckert, R., Geisel, J., and Herrmann, W. (2016). Folic acid causes higher prevalence of detectable unmetabolized folic acid in serum than B-complex: a randomized trial. Eur J Nutr 55, 1021–1028. 10.1007/s00394-015-0916-z.

12. Plumptre, L., Masih, S.P., Ly, A., Aufreiter, S., Sohn, K.J., Croxford, R., Lausman, A.Y., Berger, H., O’Connor, D.L., and Kim, Y.I. (2015). High concentrations of folate and unmetabolized folic acid in a cohort of pregnant Canadian women and umbilical cord blood. Am J Clin Nutr 102, 848–857. 10.3945/ajcn.115.110783.

13. Bailey, R.L., Mills, J.L., Yetley, E.A., Gahche, J.J., Pfeiffer, C.M., Dwyer, J.T., Dodd, K.W., Sempos, C.T., Betz, J.M., and Picciano, M.F. (2010). Unmetabolized serum folic acid and its relation to folic acid intake from diet and supplements in a nationally representative sample of adults aged > or =60 y in the United States. Am J Clin Nutr 92, 383–389. 10.3945/ajcn.2010.29499.

14. Siatka, T., Mát’uš, M., Moravcová, M., Harčárová, P., Lomozová, Z., Matoušová, K., Suwanvecho, C., Krčmová, L.K., and Mladěnka, P. (2025). Biological, dietetic and pharmacological properties of vitamin B(9). NPJ Sci Food 9, 30. 10.1038/s41538-025-00396-w.

15. Sulistyoningrum, D.C., Sullivan, T.R., Skubisz, M., Palmer, D.J., Wood, S., Ueland, P.M., McCann, A., Makrides, M., Green, T.J., and Best, K.P. (2024). Maternal serum unmetabolized folic acid concentration following multivitamin and mineral supplementation with or without folic acid after 12 weeks gestation: A randomized controlled trial. Matern Child Nutr 20, e13668. 10.1111/mcn.13668.

16. Ivanovs, A., Rybtsov, S., Ng, E.S., Stanley, E.G., Elefanty, A.G., and Medvinsky, A. (2017). Human haematopoietic stem cell development: from the embryo to the dish. Development 144, 2323–2337. 10.1242/dev.134866.

17. Dzierzak, E., and Bigas, A. (2018). Blood Development: Hematopoietic Stem Cell Dependence and Independence. Cell Stem Cell 22, 639–651. 10.1016/j.stem.2018.04.015.

18. Copp, A.J., and Greene, N.D. (2010). Genetics and development of neural tube defects. J Pathol 220, 217–230. 10.1002/path.2643.

19. Fox, J.T., and Stover, P.J. (2008). Folate-mediated one-carbon metabolism. Vitam Horm 79, 1–44. 10.1016/s0083-6729(08)00401-9.

20. Lan, X., Field, M.S., and Stover, P.J. (2018). Cell cycle regulation of folate-mediated one-carbon metabolism. Wiley Interdiscip Rev Syst Biol Med 10, e1426. 10.1002/wsbm.1426.

21. Beaudin, A.E., and Stover, P.J. (2007). Folate-mediated one-carbon metabolism and neural tube defects: balancing genome synthesis and gene expression. Birth Defects Res C Embryo Today 81, 183–203. 10.1002/bdrc.20100.

22. Ducker, G.S., and Rabinowitz, J.D. (2017). One-Carbon Metabolism in Health and Disease. Cell Metab 25, 27–42. 10.1016/j.cmet.2016.08.009.

23. Sureshchandra, S., Chan, C.N., Robino, J.J., Parmelee, L.K., Nash, M.J., Wesolowski, S.R., Pietras, E.M., Friedman, J.E., Takahashi, D., Shen, W., et al. (2022). Maternal Western-style diet remodels the transcriptional landscape of fetal hematopoietic stem and progenitor cells in rhesus macaques. Stem Cell Reports 17, 2595–2609. 10.1016/j.stemcr.2022.10.003.

24. Nash, M.J., Dobrinskikh, E., Soderborg, T.K., Janssen, R.C., Takahashi, D.L., Dean, T.A., Varlamov, O., Hennebold, J.D., Gannon, M., Aagaard, K.M., et al. (2023). Maternal diet alters long-term innate immune cell memory in fetal and juvenile hematopoietic stem and progenitor cells in nonhuman primate offspring. Cell Rep 42, 112393. 10.1016/j.celrep.2023.112393.

25. Karigane, D., Kobayashi, H., Morikawa, T., Ootomo, Y., Sakai, M., Nagamatsu, G., Kubota, Y., Goda, N., Matsumoto, M., Nishimura, E.K., et al. (2016). p38α Activates Purine Metabolism to Initiate Hematopoietic Stem/Progenitor Cell Cycling in Response to Stress. Cell Stem Cell 19, 192–204. 10.1016/j.stem.2016.05.013.

26. Orkin, S.H., and Hochedlinger, K. (2011). Chromatin connections to pluripotency and cellular reprogramming. Cell 145, 835–850. 10.1016/j.cell.2011.05.019.

27. Challen, G.A., Sun, D., Jeong, M., Luo, M., Jelinek, J., Berg, J.S., Bock, C., Vasanthakumar, A., Gu, H., Xi, Y., et al. (2011). Dnmt3a is essential for hematopoietic stem cell differentiation. Nat Genet 44, 23–31. 10.1038/ng.1009.

28. Pandolfi, A., Barreyro, L., and Steidl, U. (2013). Concise review: preleukemic stem cells: molecular biology and clinical implications of the precursors to leukemia stem cells. Stem Cells Transl Med 2, 143–150. 10.5966/sctm.2012-0109.

29. Ito, K., and Suda, T. (2014). Metabolic requirements for the maintenance of self-renewing stem cells. Nat Rev Mol Cell Biol 15, 243–256. 10.1038/nrm3772.

30. Nakamura-Ishizu, A., Ito, K., and Suda, T. (2020). Hematopoietic Stem Cell Metabolism during Development and Aging. Dev Cell 54, 239–255. 10.1016/j.devcel.2020.06.029.

31. Morganti, C., Cabezas-Wallscheid, N., and Ito, K. (2022). Metabolic Regulation of Hematopoietic Stem Cells. Hemasphere 6, e740. 10.1097/hs9.0000000000000740.

32. Takubo, K., Nagamatsu, G., Kobayashi, C.I., Nakamura-Ishizu, A., Kobayashi, H., Ikeda, E., Goda, N., Rahimi, Y., Johnson, R.S., Soga, T., et al. (2013). Regulation of glycolysis by Pdk functions as a metabolic checkpoint for cell cycle quiescence in hematopoietic stem cells. Cell Stem Cell 12, 49–61. 10.1016/j.stem.2012.10.011.

33. Takubo, K., Goda, N., Yamada, W., Iriuchishima, H., Ikeda, E., Kubota, Y., Shima, H., Johnson, R.S., Hirao, A., Suematsu, M., and Suda, T. (2010). Regulation of the HIF-1alpha level is essential for hematopoietic stem cells. Cell Stem Cell 7, 391–402. 10.1016/j.stem.2010.06.020.

34. Vannini, N., Girotra, M., Naveiras, O., Nikitin, G., Campos, V., Giger, S., Roch, A., Auwerx, J., and Lutolf, M.P. (2016). Specification of haematopoietic stem cell fate via modulation of mitochondrial activity. Nat Commun 7, 13125. 10.1038/ncomms13125.

35. Suda, T., Takubo, K., and Semenza, G.L. (2011). Metabolic regulation of hematopoietic stem cells in the hypoxic niche. Cell Stem Cell 9, 298–310. 10.1016/j.stem.2011.09.010.

36. Ho, T.T., Warr, M.R., Adelman, E.R., Lansinger, O.M., Flach, J., Verovskaya, E.V., Figueroa, M.E., and Passegué, E. (2017). Autophagy maintains the metabolism and function of young and old stem cells. Nature 543, 205–210. 10.1038/nature21388.

37. Signer, R.A., Magee, J.A., Salic, A., and Morrison, S.J. (2014). Haematopoietic stem cells require a highly regulated protein synthesis rate. Nature 509, 49–54. 10.1038/nature13035.

38. Mortensen, M., Soilleux, E.J., Djordjevic, G., Tripp, R., Lutteropp, M., Sadighi-Akha, E., Stranks, A.J., Glanville, J., Knight, S., Jacobsen, S.E., et al. (2011). The autophagy protein Atg7 is essential for hematopoietic stem cell maintenance. J Exp Med 208, 455–467. 10.1084/jem.20101145.

39. Du, C., Liu, C., Yu, K., Zhang, S., Fu, Z., Chen, X., Liao, W., Chen, J., Zhang, Y., Wang, X., et al. (2024). Mitochondrial serine catabolism safeguards maintenance of the hematopoietic stem cell pool in homeostasis and injury. Cell Stem Cell 31, 1484–1500.e1489. 10.1016/j.stem.2024.07.009.

40. LeBlanc, D.P., Behan, N.A., O’Brien, J.M., Marchetti, F., and MacFarlane, A.J. (2018). Folate deficiency increases chromosomal damage and mutations in hematopoietic cells in the transgenic mutamouse model. Environ Mol Mutagen 59, 366–374. 10.1002/em.22190.

41. Abarinov, E.V., Beaudin, A.E., Field, M.S., Perry, C.A., Allen, R.H., Stabler, S.P., and Stover, P.J. (2013). Disruption of shmt1 impairs hippocampal neurogenesis and mnemonic function in mice. J Nutr 143, 1028–1035. 10.3945/jn.113.174417.

42. MacFarlane, A.J., Behan, N.A., Field, M.S., Williams, A., Stover, P.J., and Yauk, C.L. (2015). Dietary folic acid protects against genotoxicity in the red blood cells of mice. Mutat Res 779, 105–111. 10.1016/j.mrfmmm.2015.06.012.

43. Swayne, B.G., Behan, N.A., Williams, A., Stover, P.J., Yauk, C.L., and MacFarlane, A.J. (2012). Supplemental dietary folic acid has no effect on chromosome damage in erythrocyte progenitor cells of mice. J Nutr 142, 813–817. 10.3945/jn.112.157750.

44. Benz, C., Martins, V.C., Radtke, F., and Bleul, C.C. (2008). The stream of precursors that colonizes the thymus proceeds selectively through the early T lineage precursor stage of T cell development. J Exp Med 205, 1187–1199. 10.1084/jem.20072168.

45. Muzumdar, M.D., Tasic, B., Miyamichi, K., Li, L., and Luo, L. (2007). A global double-fluorescent Cre reporter mouse. Genesis 45, 593–605. 10.1002/dvg.20335.

46. López, D.A., Apostol, A.C., Lebish, E.J., Valencia, C.H., Romero-Mulero, M.C., Pavlovich, P.V., Hernandez, G.E., Forsberg, E.C., Cabezas-Wallscheid, N., and Beaudin, A.E. (2022). Prenatal inflammation perturbs murine fetal hematopoietic development and causes persistent changes to postnatal immunity. Cell Rep 41, 111677. 10.1016/j.celrep.2022.111677.

47. Nemkov, T., Reisz, J.A., Gehrke, S., Hansen, K.C., and D’Alessandro, A. (2019). High-Throughput Metabolomics: Isocratic and Gradient Mass Spectrometry-Based Methods. Methods Mol Biol 1978, 13–26. 10.1007/978-1-4939-9236-2_2.

48. Hinge, A., He, J., Bartram, J., Javier, J., Xu, J., Fjellman, E., Sesaki, H., Li, T., Yu, J., Wunderlich, M., et al. (2020). Asymmetrically Segregated Mitochondria Provide Cellular Memory of Hematopoietic Stem Cell Replicative History and Drive HSC Attrition. Cell Stem Cell 26, 420–430.e426. 10.1016/j.stem.2020.01.016.

49. Kucinski, I., Campos, J., Barile, M., Severi, F., Bohin, N., Moreira, P.N., Allen, L., Lawson, H., Haltalli, M.L.R., Kinston, S.J., et al. (2024). A time- and single-cell-resolved model of murine bone marrow hematopoiesis. Cell Stem Cell 31, 244–259.e210. 10.1016/j.stem.2023.12.001.

50. Cabezas-Wallscheid, N., Klimmeck, D., Hansson, J., Lipka, D.B., Reyes, A., Wang, Q., Weichenhan, D., Lier, A., von Paleske, L., Renders, S., et al. (2014). Identification of regulatory networks in HSCs and their immediate progeny via integrated proteome, transcriptome, and DNA methylome analysis. Cell Stem Cell 15, 507–522. 10.1016/j.stem.2014.07.005.

51. Ewels, P.A., Peltzer, A., Fillinger, S., Patel, H., Alneberg, J., Wilm, A., Garcia, M.U., Di Tommaso, P., and Nahnsen, S. (2020). The nf-core framework for community-curated bioinformatics pipelines. Nat Biotechnol 38, 276–278. 10.1038/s41587-020-0439-x.

52. Love, M.I., Huber, W., and Anders, S. (2014). Moderated estimation of fold change and dispersion for RNA-seq data with DESeq2. Genome Biol 15, 550. 10.1186/s13059-014-0550-8.

53. Suh, J.R., Oppenheim, E.W., Girgis, S., and Stover, P.J. (2000). Purification and properties of a folate-catabolizing enzyme. J Biol Chem 275, 35646–35655. 10.1074/jbc.M005864200.

54. Bensadoun, A., and Weinstein, D. (1976). Assay of proteins in the presence of interfering materials. Anal Biochem 70, 241–250. 10.1016/s0003-2697(76)80064-4.

55. Beaudin, A.E., Boyer, S.W., Perez-Cunningham, J., Hernandez, G.E., Derderian, S.C., Jujjavarapu, C., Aaserude, E., MacKenzie, T., and Forsberg, E.C. (2016). A Transient Developmental Hematopoietic Stem Cell Gives Rise to Innate-like B and T Cells. Cell Stem Cell 19, 768–783. 10.1016/j.stem.2016.08.013.

56. Kim, E., Wright, M.J., Sioson, L., Novos, T., Gill, A.J., Benn, D.E., White, C., Dwight, T., and Clifton-Bligh, R.J. (2017). Utility of the succinate: Fumarate ratio for assessing SDH dysfunction in different tumor types. Mol Genet Metab Rep 10, 45–49. 10.1016/j.ymgmr.2016.12.006.

57. El-Gharbawy, A., and Vockley, J. (2018). Inborn Errors of Metabolism with Myopathy: Defects of Fatty Acid Oxidation and the Carnitine Shuttle System. Pediatr Clin North Am 65, 317–335. 10.1016/j.pcl.2017.11.006.

58. Dambrova, M., Makrecka-Kuka, M., Kuka, J., Vilskersts, R., Nordberg, D., Attwood, M.M., Smesny, S., Sen, Z.D., Guo, A.C., Oler, E., et al. (2022). Acylcarnitines: Nomenclature, Biomarkers, Therapeutic Potential, Drug Targets, and Clinical Trials. Pharmacol Rev 74, 506–551. 10.1124/pharmrev.121.000408.

59. Kukurugya, M.A., Rosset, S., and Titov, D.V. (2024). The Warburg Effect is the result of faster ATP production by glycolysis than respiration. Proceedings of the National Academy of Sciences 121, e2409509121. doi:10.1073/pnas.2409509121.

60. Cogliati, S., Enriquez, J.A., and Scorrano, L. (2016). Mitochondrial Cristae: Where Beauty Meets Functionality. Trends Biochem Sci 41, 261–273. 10.1016/j.tibs.2016.01.001.

61. Yu, J., Luo, Y., Lin, J., Li, Z., Fang, Z., He, H., Yan, C., and Song, Z. (2025). Mitochondrial cristae remodeling: Mechanisms, functions, and pathology. Cell Insight 4, 100285. 10.1016/j.cellin.2025.100285.

62. Anand, R., Strecker, V., Urbach, J., Wittig, I., and Reichert, A.S. (2016). Mic13 Is Essential for Formation of Crista Junctions in Mammalian Cells. PLoS One 11, e0160258. 10.1371/journal.pone.0160258.

63. Genin, E.C., Plutino, M., Bannwarth, S., Villa, E., Cisneros-Barroso, E., Roy, M., Ortega-Vila, B., Fragaki, K., Lespinasse, F., Pinero-Martos, E., et al. (2016). CHCHD10 mutations promote loss of mitochondrial cristae junctions with impaired mitochondrial genome maintenance and inhibition of apoptosis. EMBO Mol Med 8, 58–72. 10.15252/emmm.201505496.

64. Joaquim, M., Altin, S., Bulimaga, M.B., Simões, T., Nolte, H., Bader, V., Franchino, C.A., Plouzennec, S., Szczepanowska, K., Marchesan, E., et al. (2025). Mitofusin 2 displays fusion-independent roles in proteostasis surveillance. Nat Commun 16, 1501. 10.1038/s41467-025-56673-5.

65. Park, C.B., Asin-Cayuela, J., Cámara, Y., Shi, Y., Pellegrini, M., Gaspari, M., Wibom, R., Hultenby, K., Erdjument-Bromage, H., Tempst, P., et al. (2007). MTERF3 is a negative regulator of mammalian mtDNA transcription. Cell 130, 273–285. 10.1016/j.cell.2007.05.046.

66. Cámara, Y., Asin-Cayuela, J., Park, C.B., Metodiev, M.D., Shi, Y., Ruzzenente, B., Kukat, C., Habermann, B., Wibom, R., Hultenby, K., et al. (2011). MTERF4 regulates translation by targeting the methyltransferase NSUN4 to the mammalian mitochondrial ribosome. Cell Metab 13, 527–539. 10.1016/j.cmet.2011.04.002.

67. Stiburek, L., Cesnekova, J., Kostkova, O., Fornuskova, D., Vinsova, K., Wenchich, L., Houstek, J., and Zeman, J. (2012). YME1L controls the accumulation of respiratory chain subunits and is required for apoptotic resistance, cristae morphogenesis, and cell proliferation. Mol Biol Cell 23, 1010–1023. 10.1091/mbc.E11-08-0674.

68. Hidalgo San Jose, L., and Signer, R.A.J. (2019). Cell-type-specific quantification of protein synthesis in vivo. Nat Protoc 14, 441–460. 10.1038/s41596-018-0100-z.

69. Chen, M.Z., Moily, N.S., Bridgford, J.L., Wood, R.J., Radwan, M., Smith, T.A., Song, Z., Tang, B.Z., Tilley, L., Xu, X., et al. (2017). A thiol probe for measuring unfolded protein load and proteostasis in cells. Nat Commun 8, 474. 10.1038/s41467-017-00203-5.

70. Aguilo, F., Avagyan, S., Labar, A., Sevilla, A., Lee, D.F., Kumar, P., Lemischka, I.R., Zhou, B.Y., and Snoeck, H.W. (2011). Prdm16 is a physiologic regulator of hematopoietic stem cells. Blood 117, 5057–5066. 10.1182/blood-2010-08-300145.

71. Gudmundsson, K.O., Nguyen, N., Oakley, K., Han, Y., Gudmundsdottir, B., Liu, P., Tessarollo, L., Jenkins, N.A., Copeland, N.G., and Du, Y. (2020). Prdm16 is a critical regulator of adult long-term hematopoietic stem cell quiescence. Proc Natl Acad Sci U S A 117, 31945–31953. 10.1073/pnas.2017626117.

72. Chen, J.Y., Miyanishi, M., Wang, S.K., Yamazaki, S., Sinha, R., Kao, K.S., Seita, J., Sahoo, D., Nakauchi, H., and Weissman, I.L. (2016). Hoxb5 marks long-term haematopoietic stem cells and reveals a homogenous perivascular niche. Nature 530, 223–227. 10.1038/nature16943.

73. Huo, Y., Li, B.Y., Lin, Z.F., Wang, W., Jiang, X.X., Chen, X., Xi, W.J., Yang, A.G., Chen, S.Y., and Wang, T. (2018). MYSM1 Is Essential for Maintaining Hematopoietic Stem Cell (HSC) Quiescence and Survival. Med Sci Monit 24, 2541–2549. 10.12659/msm.906876.

74. Sullivan, L.B., Luengo, A., Danai, L.V., Bush, L.N., Diehl, F.F., Hosios, A.M., Lau, A.N., Elmiligy, S., Malstrom, S., Lewis, C.A., and Vander Heiden, M.G. (2018). Aspartate is an endogenous metabolic limitation for tumour growth. Nat Cell Biol 20, 782–788. 10.1038/s41556-018-0125-0.

75. Westermann, B. (2010). Mitochondrial fusion and fission in cell life and death. Nat Rev Mol Cell Biol 11, 872–884. 10.1038/nrm3013.

76. Lu, Q., Wang, P.S., and Yang, L. (2021). Golgi-associated Rab GTPases implicated in autophagy. Cell Biosci 11, 35. 10.1186/s13578-021-00543-2.

77. Morgan, N.E., Cutrona, M.B., and Simpson, J.C. (2019). Multitasking Rab Proteins in Autophagy and Membrane Trafficking: A Focus on Rab33b. Int J Mol Sci 20. 10.3390/ijms20163916.

78. Ferreira, A., Pereira, F., Reis, C., Oliveira, M.J., Sousa, M.J., and Preto, A. (2022). Crucial Role of Oncogenic KRAS Mutations in Apoptosis and Autophagy Regulation: Therapeutic Implications. Cells 11. 10.3390/cells11142183.

79. Fry, M.Y., Navarro, P.P., Hakim, P., Ananda, V.Y., Qin, X., Landoni, J.C., Rath, S., Inde, Z., Lugo, C.M., Luce, B.E., et al. (2024). In situ architecture of Opa1-dependent mitochondrial cristae remodeling. Embo j 43, 391–413. 10.1038/s44318-024-00027-2.

80. Halling, J.F., and Pilegaard, H. (2020). PGC-1α-mediated regulation of mitochondrial function and physiological implications. Appl Physiol Nutr Metab 45, 927–936. 10.1139/apnm-2020-0005.

81. Frezza, C., Cipolat, S., Martins de Brito, O., Micaroni, M., Beznoussenko, G.V., Rudka, T., Bartoli, D., Polishuck, R.S., Danial, N.N., De Strooper, B., and Scorrano, L. (2006). OPA1 controls apoptotic cristae remodeling independently from mitochondrial fusion. Cell 126, 177–189. 10.1016/j.cell.2006.06.025.

82. Van Humbeeck, C., Cornelissen, T., and Vandenberghe, W. (2011). Ambra1: a Parkin-binding protein involved in mitophagy. Autophagy 7, 1555–1556. 10.4161/auto.7.12.17893.

83. Strappazzon, F., Nazio, F., Corrado, M., Cianfanelli, V., Romagnoli, A., Fimia, G.M., Campello, S., Nardacci, R., Piacentini, M., Campanella, M., and Cecconi, F. (2015). AMBRA1 is able to induce mitophagy via LC3 binding, regardless of PARKIN and p62/SQSTM1. Cell Death Differ 22, 419–432. 10.1038/cdd.2014.139.

84. Liang, C., Lee, J.S., Inn, K.S., Gack, M.U., Li, Q., Roberts, E.A., Vergne, I., Deretic, V., Feng, P., Akazawa, C., and Jung, J.U. (2008). Beclin1-binding UVRAG targets the class C Vps complex to coordinate autophagosome maturation and endocytic trafficking. Nat Cell Biol 10, 776–787. 10.1038/ncb1740.

85. Issa, A.R., Sun, J., Petitgas, C., Mesquita, A., Dulac, A., Robin, M., Mollereau, B., Jenny, A., Chérif-Zahar, B., and Birman, S. (2018). The lysosomal membrane protein LAMP2A promotes autophagic flux and prevents SNCA-induced Parkinson disease-like symptoms in the Drosophila brain. Autophagy 14, 1898–1910. 10.1080/15548627.2018.1491489.

86. Jaber, N., Dou, Z., Chen, J.S., Catanzaro, J., Jiang, Y.P., Ballou, L.M., Selinger, E., Ouyang, X., Lin, R.Z., Zhang, J., and Zong, W.X. (2012). Class III PI3K Vps34 plays an essential role in autophagy and in heart and liver function. Proc Natl Acad Sci U S A 109, 2003–2008. 10.1073/pnas.1112848109.

87. Takahashi, Y., Meyerkord, C.L., Hori, T., Runkle, K., Fox, T.E., Kester, M., Loughran, T.P., and Wang, H.G. (2011). Bif-1 regulates Atg9 trafficking by mediating the fission of Golgi membranes during autophagy. Autophagy 7, 61–73. 10.4161/auto.7.1.14015.

88. Madan, V., Han, L., Hattori, N., Teoh, W.W., Mayakonda, A., Sun, Q.Y., Ding, L.W., Nordin, H.B.M., Lim, S.L., Shyamsunder, P., et al. (2018). ASXL2 regulates hematopoiesis in mice and its deficiency promotes myeloid expansion. Haematologica 103, 1980–1990. 10.3324/haematol.2018.189928.

89. Antunes, E.T.B., and Ottersbach, K. (2020). The MLL/SET family and haematopoiesis. Biochim Biophys Acta Gene Regul Mech 1863, 194579. 10.1016/j.bbagrm.2020.194579.

90. Lin, C.C., Yao, C.Y., Hsu, Y.C., Hou, H.A., Yuan, C.T., Li, Y.H., Kao, C.J., Chuang, P.H., Chiu, Y.C., Chen, Y., et al. (2020). Knock-out of Hopx disrupts stemness and quiescence of hematopoietic stem cells in mice. Oncogene 39, 5112–5123. 10.1038/s41388-020-1340-2.

91. Florian, M.C., Dörr, K., Niebel, A., Daria, D., Schrezenmeier, H., Rojewski, M., Filippi, M.D., Hasenberg, A., Gunzer, M., Scharffetter-Kochanek, K., et al. (2012). Cdc42 activity regulates hematopoietic stem cell aging and rejuvenation. Cell Stem Cell 10, 520–530. 10.1016/j.stem.2012.04.007.

92. Reid, T., Bathoorn, A., Ahmadian, M.R., and Collard, J.G. (1999). Identification and characterization of hPEM-2, a guanine nucleotide exchange factor specific for Cdc42. J Biol Chem 274, 33587–33593. 10.1074/jbc.274.47.33587.

93. Cotteret, S., Jaffer, Z.M., Beeser, A., and Chernoff, J. (2003). p21-Activated kinase 5 (Pak5) localizes to mitochondria and inhibits apoptosis by phosphorylating BAD. Mol Cell Biol 23, 5526–5539. 10.1128/mcb.23.16.5526-5539.2003.

94. Sumi, T., Matsumoto, K., Takai, Y., and Nakamura, T. (1999). Cofilin phosphorylation and actin cytoskeletal dynamics regulated by rho- and Cdc42-activated LIM-kinase 2. J Cell Biol 147, 1519–1532. 10.1083/jcb.147.7.1519.

95. Ohta, Y., Suzuki, N., Nakamura, S., Hartwig, J.H., and Stossel, T.P. (1999). The small GTPase RalA targets filamin to induce filopodia. Proc Natl Acad Sci U S A 96, 2122–2128. 10.1073/pnas.96.5.2122.

96. Mak, K.S., Funnell, A.P., Pearson, R.C., and Crossley, M. (2011). PU.1 and Haematopoietic Cell Fate: Dosage Matters. Int J Cell Biol 2011, 808524. 10.1155/2011/808524.

97. Yokota, H., Goldring, M.B., and Sun, H.B. (2003). CITED2-mediated regulation of MMP-1 and MMP-13 in human chondrocytes under flow shear. J Biol Chem 278, 47275–47280. 10.1074/jbc.M304652200.

98. Elvert, G., Kappel, A., Heidenreich, R., Englmeier, U., Lanz, S., Acker, T., Rauter, M., Plate, K., Sieweke, M., Breier, G., and Flamme, I. (2003). Cooperative interaction of hypoxia-inducible factor-2alpha (HIF-2alpha) and Ets-1 in the transcriptional activation of vascular endothelial growth factor receptor-2 (Flk-1). J Biol Chem 278, 7520–7530. 10.1074/jbc.M211298200.

99. Chowdhury, I., Mo, Y., Gao, L., Kazi, A., Fisher, A.B., and Feinstein, S.I. (2009). Oxidant stress stimulates expression of the human peroxiredoxin 6 gene by a transcriptional mechanism involving an antioxidant response element. Free Radic Biol Med 46, 146–153. 10.1016/j.freeradbiomed.2008.09.027.

100. Liu, Y., Adachi, M., Zhao, S., Hareyama, M., Koong, A.C., Luo, D., Rando, T.A., Imai, K., and Shinomura, Y. (2009). Preventing oxidative stress: a new role for XBP1. Cell Death Differ 16, 847–857. 10.1038/cdd.2009.14.

101. Liu, L., Zhao, M., Jin, X., Ney, G., Yang, K.B., Peng, F., Cao, J., Iwawaki, T., Del Valle, J., Chen, X., and Li, Q. (2019). Adaptive endoplasmic reticulum stress signalling via IRE1α-XBP1 preserves self-renewal of haematopoietic and pre-leukaemic stem cells. Nat Cell Biol 21, 328–337. 10.1038/s41556-019-0285-6.

102. Knudsen, K.J., Rehn, M., Hasemann, M.S., Rapin, N., Bagger, F.O., Ohlsson, E., Willer, A., Frank, A.K., Søndergaard, E., Jendholm, J., et al. (2015). ERG promotes the maintenance of hematopoietic stem cells by restricting their differentiation. Genes Dev 29, 1915–1929. 10.1101/gad.268409.115.

103. Ng, A.P., Loughran, S.J., Metcalf, D., Hyland, C.D., de Graaf, C.A., Hu, Y., Smyth, G.K., Hilton, D.J., Kile, B.T., and Alexander, W.S. (2011). Erg is required for self-renewal of hematopoietic stem cells during stress hematopoiesis in mice. Blood 118, 2454–2461. 10.1182/blood-2011-03-344739.

104. Bracken, A.P., Kleine-Kohlbrecher, D., Dietrich, N., Pasini, D., Gargiulo, G., Beekman, C., Theilgaard-Mönch, K., Minucci, S., Porse, B.T., Marine, J.C., et al. (2007). The Polycomb group proteins bind throughout the INK4A-ARF locus and are disassociated in senescent cells. Genes Dev 21, 525–530. 10.1101/gad.415507.

105. Zhang, J., Bonasio, R., Strino, F., Kluger, Y., Holloway, J.K., Modzelewski, A.J., Cohen, P.E., and Reinberg, D. (2013). SFMBT1 functions with LSD1 to regulate expression of canonical histone genes and chromatin-related factors. Genes Dev 27, 749–766. 10.1101/gad.210963.112.

106. Tavares, L., Dimitrova, E., Oxley, D., Webster, J., Poot, R., Demmers, J., Bezstarosti, K., Taylor, S., Ura, H., Koide, H., et al. (2012). RYBP-PRC1 complexes mediate H2A ubiquitylation at polycomb target sites independently of PRC2 and H3K27me3. Cell 148, 664–678. 10.1016/j.cell.2011.12.029.

107. Domen, J., Cheshier, S.H., and Weissman, I.L. (2000). The role of apoptosis in the regulation of hematopoietic stem cells: Overexpression of Bcl-2 increases both their number and repopulation potential. J Exp Med 191, 253–264. 10.1084/jem.191.2.253.

108. Yang, J., Aguila, J.R., Alipio, Z., Lai, R., Fink, L.M., and Ma, Y. (2011). Enhanced self-renewal of hematopoietic stem/progenitor cells mediated by the stem cell gene Sall4. J Hematol Oncol 4, 38. 10.1186/1756-8722-4-38.

109. Zhang, S., Hulver, M.W., McMillan, R.P., Cline, M.A., and Gilbert, E.R. (2014). The pivotal role of pyruvate dehydrogenase kinases in metabolic flexibility. Nutr Metab (Lond) 11, 10. 10.1186/1743-7075-11-10.

110. Wilson, J.E. (2003). Isozymes of mammalian hexokinase: structure, subcellular localization and metabolic function. J Exp Biol 206, 2049–2057. 10.1242/jeb.00241.

111. Gabriele, M., Brandão, H.B., Grosse-Holz, S., Jha, A., Dailey, G.M., Cattoglio, C., Hsieh, T.S., Mirny, L., Zechner, C., and Hansen, A.S. (2022). Dynamics of CTCF- and cohesin-mediated chromatin looping revealed by live-cell imaging. Science 376, 496–501. 10.1126/science.abn6583.

112. Kim, S., Yu, N.K., and Kaang, B.K. (2015). CTCF as a multifunctional protein in genome regulation and gene expression. Exp Mol Med 47, e166. 10.1038/emm.2015.33.

113. Qi, Q., Cheng, L., Tang, X., He, Y., Li, Y., Yee, T., Shrestha, D., Feng, R., Xu, P., Zhou, X., et al. (2021). Dynamic CTCF binding directly mediates interactions among cis-regulatory elements essential for hematopoiesis. Blood 137, 1327–1339. 10.1182/blood.2020005780.

114. Qiu, Y., and Huang, S. (2020). CTCF-mediated genome organization and leukemogenesis. Leukemia 34, 2295–2304. 10.1038/s41375-020-0906-x.

115. Kim, T.G., Kim, S., Jung, S., Kim, M., Yang, B., Lee, M.G., and Kim, H.P. (2017). CCCTC-binding factor is essential to the maintenance and quiescence of hematopoietic stem cells in mice. Exp Mol Med 49, e371. 10.1038/emm.2017.124.

116. López, D.A., Otsuka, K.S., Apostol, A.C., Posada, J., Sánchez-Arcila, J.C., Jensen, K.D., and Beaudin, A.E. (2023). Both maternal IFNγ exposure and acute prenatal infection with Toxoplasma gondii activate fetal hematopoietic stem cells. Embo j 42, e112693. 10.15252/embj.2022112693.

117. Cabezas-Wallscheid, N., Buettner, F., Sommerkamp, P., Klimmeck, D., Ladel, L., Thalheimer, F.B., Pastor-Flores, D., Roma, L.P., Renders, S., Zeisberger, P., et al. (2017). Vitamin A-Retinoic Acid Signaling Regulates Hematopoietic Stem Cell Dormancy. Cell 169, 807–823.e819. 10.1016/j.cell.2017.04.018.

118. Chen, C., Yu, W., Tober, J., Gao, P., He, B., Lee, K., Trieu, T., Blobel, G.A., Speck, N.A., and Tan, K. (2019). Spatial Genome Re-organization between Fetal and Adult Hematopoietic Stem Cells. Cell Rep 29, 4200–4211.e4207. 10.1016/j.celrep.2019.11.065.

119. Fan, J., Ye, J., Kamphorst, J.J., Shlomi, T., Thompson, C.B., and Rabinowitz, J.D. (2014). Quantitative flux analysis reveals folate-dependent NADPH production. Nature 510, 298–302. 10.1038/nature13236.

120. Scudese, E., Marshall, A.G., Vue, Z., Exil, V., Rodriguez, B.I., Demirci, M., Vang, L., López, E.G., Neikirk, K., Shao, B., et al. (2025). 3D Mitochondrial Structure in Aging Human Skeletal Muscle: Insights Into MFN-2-Mediated Changes. Aging Cell 24, e70054. 10.1111/acel.70054.

121. Luchsinger, L.L., de Almeida, M.J., Corrigan, D.J., Mumau, M., and Snoeck, H.W. (2016). Mitofusin 2 maintains haematopoietic stem cells with extensive lymphoid potential. Nature 529, 528–531. 10.1038/nature16500.

122. Farhan, H., and Hsu, V.W. (2016). Cdc42 and Cellular Polarity: Emerging Roles at the Golgi. Trends Cell Biol 26, 241–248. 10.1016/j.tcb.2015.11.003.

123. Martinez-Lopez, N., Mattar, P., Toledo, M., Bains, H., Kalyani, M., Aoun, M.L., Sharma, M., McIntire, L.B.J., Gunther-Cummins, L., Macaluso, F.P., et al. (2023). mTORC2-NDRG1-CDC42 axis couples fasting to mitochondrial fission. Nat Cell Biol 25, 989–1003. 10.1038/s41556-023-01163-3.

124. Wellen, K.E., Hatzivassiliou, G., Sachdeva, U.M., Bui, T.V., Cross, J.R., and Thompson, C.B. (2009). ATP-citrate lyase links cellular metabolism to histone acetylation. Science 324, 1076–1080. 10.1126/science.1164097.

125. Kaelin, W.G., Jr., and McKnight, S.L. (2013). Influence of metabolism on epigenetics and disease. Cell 153, 56–69. 10.1016/j.cell.2013.03.004.

126. Reid, M.A., Dai, Z., and Locasale, J.W. (2017). The impact of cellular metabolism on chromatin dynamics and epigenetics. Nat Cell Biol 19, 1298–1306. 10.1038/ncb3629.

127. Lourenço Dos Santos, S., Petropoulos, I., and Friguet, B. (2018). The Oxidized Protein Repair Enzymes Methionine Sulfoxide Reductases and Their Roles in Protecting against Oxidative Stress, in Ageing and in Regulating Protein Function. Antioxidants (Basel) 7. 10.3390/antiox7120191.

128. Li, Y., He, Y., Zheng, Q., Zhang, J., Pan, X., Zhang, X., Yuan, H., Wang, G., Liu, X., Zhou, X., et al. (2025). Mitochondrial pyruvate carriers control airway basal progenitor cell function through glycolytic-epigenetic reprogramming. Cell Stem Cell 32, 105–120.e106. 10.1016/j.stem.2024.09.015.

129. Sutendra, G., Kinnaird, A., Dromparis, P., Paulin, R., Stenson, T.H., Haromy, A., Hashimoto, K., Zhang, N., Flaim, E., and Michelakis, E.D. (2014). A nuclear pyruvate dehydrogenase complex is important for the generation of acetyl-CoA and histone acetylation. Cell 158, 84–97. 10.1016/j.cell.2014.04.046.

130. Chua, B.A., and Signer, R.A.J. (2020). Hematopoietic stem cell regulation by the proteostasis network. Curr Opin Hematol 27, 254–263. 10.1097/moh.0000000000000591.

131. Konuma, T., Oguro, H., and Iwama, A. (2010). Role of the polycomb group proteins in hematopoietic stem cells. Dev Growth Differ 52, 505–516. 10.1111/j.1440-169X.2010.01191.x.

132. Klauke, K., and de Haan, G. (2011). Polycomb group proteins in hematopoietic stem cell aging and malignancies. Int J Hematol 94, 11–23. 10.1007/s12185-011-0857-0.

133. Shyh-Chang, N., Daley, G.Q., and Cantley, L.C. (2013). Stem cell metabolism in tissue development and aging. Development 140, 2535–2547. 10.1242/dev.091777.

134. Ly, C.H., Lynch, G.S., and Ryall, J.G. (2020). A Metabolic Roadmap for Somatic Stem Cell Fate. Cell Metab 31, 1052–1067. 10.1016/j.cmet.2020.04.022.

135. Henry, C.J., Nemkov, T., Casás-Selves, M., Bilousova, G., Zaberezhnyy, V., Higa, K.C., Serkova, N.J., Hansen, K.C., D’Alessandro, A., and DeGregori, J. (2017). Folate dietary insufficiency and folic acid supplementation similarly impair metabolism and compromise hematopoiesis. Haematologica 102, 1985–1994. 10.3324/haematol.2017.171074.

136. Mason, J.B., Dickstein, A., Jacques, P.F., Haggarty, P., Selhub, J., Dallal, G., and Rosenberg, I.H. (2007). A temporal association between folic acid fortification and an increase in colorectal cancer rates may be illuminating important biological principles: a hypothesis. Cancer Epidemiol Biomarkers Prev 16, 1325–1329. 10.1158/1055-9965.Epi-07-0329.

137. Figueiredo, J.C., Grau, M.V., Haile, R.W., Sandler, R.S., Summers, R.W., Bresalier, R.S., Burke, C.A., McKeown-Eyssen, G.E., and Baron, J.A. (2009). Folic acid and risk of prostate cancer: results from a randomized clinical trial. J Natl Cancer Inst 101, 432–435. 10.1093/jnci/djp019.

138. Hirsch, S., Sanchez, H., Albala, C., de la Maza, M.P., Barrera, G., Leiva, L., and Bunout, D. (2009). Colon cancer in Chile before and after the start of the flour fortification program with folic acid. Eur J Gastroenterol Hepatol 21, 436–439. 10.1097/MEG.0b013e328306ccdb.

139. Wien, T.N., Pike, E., Wisløff, T., Staff, A., Smeland, S., and Klemp, M. (2012). Cancer risk with folic acid supplements: a systematic review and meta-analysis. BMJ Open 2, e000653. 10.1136/bmjopen-2011-000653.

140. Yoo, S., Montazeri, A., Bennett, D., Bo, Y., Chen, P., Duthie, S., Jensen, N., Kaminga, A., Lai, J.S., Li, X., et al. (2026). Folate and global health umbrella review series, part 2: syntheses on cancers. J Glob Health 16, 04102. 10.7189/jogh.16.04102.

141. Zhao, L., Zhang, Y., Liu, J., Hébert, J.R., Giovannucci, E., Zhang, X., and Steck, S.E. (2025). Trends in dietary supplement use among U.S. adults between 2011 and 2023. Eur J Nutr 64, 304. 10.1007/s00394-025-03825-4.

142. Lam, C.S., O’Connell, K., Monroy-Iglesias, M.J., Wang, P., Hou, Y.-N., Du, M., Giovannucci, E.L., Mao, J.J., and Kantor, E.D. (2026). Emerging Patterns in Dietary Supplement Use Among US Adults, 1999-2023. JAMA Network Open 9, e2619291–e2619291. 10.1001/jamanetworkopen.2026.19291.

