## Supplementary material for "Maternal Folate Nutrition Programs Adult Hematopoietic Stem Cell Function Through Mitochondrial Remodeling": All Supplemental Materials

#### SUPPLEMENTAL INFORMATION

##### Supplemental Methods

##### Mice and dietary interventions

All animal experiments were conducted in accordance with protocols approved by the University of Utah Institutional Animal Care and Use committee. C57BL/6J mice were purchased from Jax. Additional strains included Flk2-Cre [1] mice crossed with mT/mG reporter mice [2] (FlkSwitch), and congenic CD45.1 PepBoyJ mice used as transplant recipients or competitive bone marrow donors where indicated.

To manipulate maternal folate exposure, female mice were placed on experimental diets at weaning (3 weeks of age) and maintained on diets for three weeks prior to mating, throughout pregnancy, and during lactation. Diets consisted of modified AIN-93G formulations (Dyets Inc, Bethlehem, PA) containing either no folic acid (0mg/kg; folate-deficient), a control diet (2mg/kg folic acid; folate-control), or supplemented (8mg/kg folic acid; folate-supplemented). All offspring were weaned at three weeks of age onto standard Teklad 2920x chow for the remainder of the study. Adult offspring were analyzed at 8-12 weeks of age.

##### Hematopoietic stem cell analysis and cell sorting

Bone marrow was harvested from femurs, tibia and pelvic bones by mechanical dissociation and filtration to generate single-cell suspensions. Red blood cells were removed using ammonium chloride-based lysis buffer. Hematopoietic stem cells (HSCs) were identified by flow cytometry using established surface markers. Briefly, cells were first gated on $CD45^+$ hematopoietic cells and lineage-negative populations (Lin-), defined by absence of staining for CD3, CD4, CD5, CD8, CD19, Ter119 and CD11b. Within the Lin- compartment, HSCs were defined as $Sca1^+$, $cKit^+$, $Flk2^-$, $CD48^-$, $CD150^+$ cells. Developmentally-restricted HSCs (drHSCs) were further defined in FlkSwitch mice as $GFP^+$ cells within the HSC gate. Cell sorting and analysis were performed using fluorescence-activated cell sorting instruments, Cytek Aurora (Cytek Biosciences, Fremont, CA, USA) and BD FACSAria (BD Biosciences, Ashland, OR, USA), and data were analyzed using FlowJo software (BD Biosciences, Ashland, OR, USA).

##### Complete blood counts (CBC)

Peripheral blood was collected from fetal, neonatal, or adult mice into heparin-coated blood collection tubes. Complete blood counts were performed using an automated hematology analyzer (Drew Scientific Hemavet 950LV). Data were analyzed to assess standard hematologic parameters, including red blood cell, white blood cell, and platelet counts.

**Hematopoietic stem cell transplantation**

For competitive bone marrow transplantation assays, 200 double-sorted purified HSCs were isolated from FlkSwitch adult offspring group by flow cytometry and transplanted alongside 5 x $10^5$ whole bone marrow cells from CD45.2 wild-type mice via the retro-orbital plexus. Recipient CD45.1 PepBoyJ mice (males, 8-12 weeks of age) were lethally irradiated with two doses of 400 rads (total 8 Gy) by X-ray radiation (Rad Source RS-2000), administered 4 hours apart. Mice were maintained on 1 g/L ampicillin in the drinking water for 4 weeks post-transplant. Peripheral blood was collected from recipients monthly for 20 weeks via retro-orbital sampling (50 µL per time point) to assess donor chimerism and hematopoietic reconstitution. Chimerism and lineage contributions were determined by flow cytometry using CD45.1/CD45.2 markers and lineage-specific antibodies.

**Assessment of mitochondrial functional and lipid staining by flow cytometry**

Mitochondrial membrane potential, mitochondrial content, and neutral lipid accumulation were assessed in hematopoietic stem and progenitor cells (HSPCs) by flow cytometry using tetramethylrhodamine methyl ester (TMRM), MitoTracker Green, and BODIPY 493/503 staining, respectively. Freshly isolated bone marrow cells were first stained with fluorophore-conjugated surface antibodies as described above. Following surface staining, cells were washed and resuspended in pre-warmed staining buffer. For measurement of mitochondrial membrane potential, cells were incubated with TMRM (20 nM) for 30 minutes at 37°C in the dark. Mitochondrial mass was assessed in parallel using MitoTracker Green (20 nM) under the same conditions. Neutral lipid content was measured using BODIPY 493/503 (2 µM). Following staining, cells were washed, resuspended in ice-cold sorting buffer, and immediately analyzed. Dead cells were excluded using propidium iodide (PI) or DAPI. Flow cytometric analysis was performed on a Cytek Aurora, and fluorescence intensity was quantified within defined HSPC and HSC populations based on established gating strategies. Median fluorescence intensity (MFI) was used as a measure of relative signal for each probe. Fluorescence signals were normalized to an internal wild-type control sample included in each experiment.

**Metabolomics analysis**

**Sample preparation**

Sorted cells were counted, pelleted at 1200 rpm, and stored at −80°C until analysis. Each biological replicate consisted of pooled cells from three mice derived from independent litters, with three independent samples analyzed per experimental group. Prior to LC–MS analysis, samples were thawed on ice and resuspended in 200 µL of pre-chilled (−20°C) methanol:acetonitrile:water (5:3:2, v/v/v). Samples were vortexed continuously for 30 minutes at 4°C to facilitate metabolite extraction. Insoluble material was removed by centrifugation at 10,000 × g for 10 minutes at 4°C, and supernatants were collected and dried under vacuum (Labconco, Kansas City, KS, USA). Dried extracts were resuspended in 20 µL of 0.1% formic acid prior to analysis.

**UHPLC–MS analysis**

Untargeted metabolomic analyses were performed as previously described.[3] Briefly, samples were analyzed using a Vanquish UHPLC system coupled online to a Q Exactive mass spectrometer (Thermo Fisher Scientific, San Jose, CA, USA). Metabolites were separated on a Kinetex C18 column (2.1 × 150 mm, 1.7 µm particle size; Phenomenex, Torrance, CA, USA) equipped with a guard column (SecurityGuard™ UHPLC C18, Phenomenex). For positive ion mode, the mobile phases consisted of (A) water with 0.1% formic acid and (B) acetonitrile with 0.1% formic acid. For negative ion mode, mobile phases consisted of (A) water:acetonitrile (95:5) with 1 mM ammonium acetate and (B) acetonitrile:water (95:5) with 1 mM ammonium acetate. Samples were resolved using either isocratic elution (5% B at 250 µL/min, 25°C) or a gradient from 5% to 95% B over 1 minute followed by a 2-minute hold at 95% B (400 µL/min, 45°C). The mass spectrometer was operated independently in positive and negative ion modes, scanning in full MS mode (2 µscans) over a mass range of 60–900 m/z at 70,000 resolution. Source parameters included a spray voltage of 4 kV, sheath gas at 45, and auxiliary gas at 15. Instrument calibration was performed prior to analysis using Pierce™ Positive and Negative Ion Calibration Solutions (Thermo Fisher Scientific). Raw data files were converted to mzXML format using Mass Matrix (Cleveland, OH, USA). Samples were analyzed in randomized order, with a pooled technical mixture injected every 15 samples to monitor instrument performance.

**Data processing and analysis**

Metabolite identification, isotopologue distribution analysis, and correction for natural isotope abundance (^2H, ^13C, ^15N) were performed using MAVEN (Princeton, NJ, USA). Metabolite abundances were normalized to input cell number prior to downstream analysis. Statistical analyses, including univariate testing (Student's t-test or ANOVA), multivariate analysis (PLS-DA), metabolic pathway enrichment, and data visualization (heatmaps), were conducted using MetaboAnalyst 6.0. Hierarchical clustering analysis was additionally performed using GENE-E (Broad Institute, Cambridge, MA, USA). Graphs were generated using GraphPad Prism 5.0 (GraphPad Software Inc., La Jolla, CA, USA).

**Transmission electron microscopy**

For ultrastructural analysis, hematopoietic stem cells (HSCs) were isolated by flow cytometry and processed using a protocol adapted from Filippi, 2016.[4] Briefly, isolated cells were fixed in 1% paraformaldehyde (PFA) and 2.5% glutaraldehyde in 0.1 M sodium cacodylate buffer (pH 7.4) at 4 °C. After primary fixation, samples were washed and stained in 200 µL of a 2 mg/mL Evans Blue solution prepared in cacodylate buffer for 20 minutes at room temperature. Cells were then washed, resuspended in 200 µL of buffer in a 0.5 mL tube, and mixed with 200 µL of 4% low-melting-point agarose in cacodylate buffer. The suspension was immediately centrifuged at 1,000 × g for 10 minutes at 30 °C. Following centrifugation, the tube containing the semi-solid agarose pellet was transferred to 4 °C to allow complete solidification. The solidified pellet was carefully trimmed and transferred to a 1.5 mL tube for washing. Samples were then post-fixed in 1% osmium tetroxide, dehydrated through a graded ethanol series (25%, 50%, 75%, 95%, and 100% ethanol; 15 minutes each at room temperature), and embedded in epoxy resin (Epon 812) using a pyramid-tip rubber mold. Resin blocks were polymerized at 60 °C for 72 hours. Ultrathin sections (70-90 nm) were cut with an ultramicrotome, mounted on copper grids, stained with uranyl acetate and lead citrate, and imaged at the University of Utah Electron Microscopy Core using a JEOL JEM-1400 transmission electron microscope (JEOL USA, Inc., Peabody, MA, USA). Mitochondrial ultrastructure in HSCs was assessed based on established morphological criteria, including cristae organization and matrix density. Representative images were acquired at multiple magnifications. For quantitative analysis, mitochondrial morphology was evaluated in a blinded manner using ImageJ (National Institutes of Health, Bethesda, MD, USA). Cristae integrity was quantified by measuring the proportion of mitochondrial area lacking clearly defined cristae. Approximately 12 cells per group were analyzed across three independent biological replicates.

**Extracellular flux analysis**

Mitochondrial respiration and glycolytic function were assessed using Seahorse extracellular flux analysis (Agilent Technologies, Santa Clara, CA, USA). Fetal hematopoietic stem and progenitor cells (HSPCs) were isolated by flow cytometry and plated at 10,000 cells per well in Seahorse assay plates. Mitochondrial function was evaluated using the Mito Stress Test, and glycolytic function was assessed using the Glycolysis Stress Test, performed according to the manufacturer's instructions (Oligomycin 1 μM, FCCP 4 μM, Rotenone/Antimycin A 1 μM; Agilent Seahorse XF Analyzer (Agilent Technologies, Santa Clara, CA, USA). Each experimental group consisted of three independent biological replicates derived from separate litters, with 4 technical triplicates plated for each sample. Oxygen consumption rate (OCR) and extracellular acidification rate (ECAR) measurements were acquired and analyzed using Seahorse Wave software (Agilent Technologies, Santa Clara, CA, USA). Data were normalized to input cell number and are presented as rates per 1,000 cells.

**Immunocytochemistry**

Hematopoietic stem cells (HSCs) were isolated by flow cytometry and fixed in 4% paraformaldehyde at room temperature. Fixed cells were washed and transferred to polystyrene chamber slides mounted on glass coverslips. The chamber slides were centrifuged at 1,000 RCF for 10 minutes and then incubated at 4 °C overnight to allow complete cell adhesion. After a brief wash with PBS, cells were permeabilized with 0.2% Triton X-100 in PBS for 10 minutes and subsequently blocked in blocking buffer containing 2.5% normal goat serum and 2.5% donkey serum in PBST (PBS supplemented with 0.2% Triton X-100). Cells were then incubated with primary antibodies overnight at 4 °C: rat anti-TOMM20 (1:250), mouse anti-CDC42 (1:50), and rabbit anti-LAMP1 (1:500) in 1:1 mixture of the blocking buffer and PBST. Following washing, cells were incubated for 1 hour at room temperature with the appropriate secondary antibodies, including Alexa Fluor 488 AffiniPure donkey anti-rabbit IgG (H+L) (1:500), Cy3 AffiniPure donkey anti-rat IgG (H+L) (1:500), Alexa Fluor 488 AffiniPure donkey anti-mouse IgG (H+L) (1:500), and Hoechst 33342 (1:20,000). Cells were then washed with PBS, and coverslips were mounted using ProLong Diamond Antifade Mountant for imaging.

**Super-resolution confocal imaging and 3D reconstruction**

Super-resolution confocal imaging was performed using a Zeiss LSM 880 Airyscan microscope (Carl Zeiss Microscopy LLC, White Plains, NY, USA) at the University of Utah Fluorescence Microscopy Core Facility. Images were acquired with a 63× high-numerical-aperture objective, and Z-stacks were collected at 0.3 μm step intervals to capture the full three-dimensional architecture of mitochondria and associated subcellular structures. Image stacks were processed by Airyscan deconvolution and further analyzed using Imaris software (Bitplane AG,

Zurich, Switzerland) to normalize signal intensity throughout the Z-stack. Three-dimensional reconstructions were generated for quantitative analysis of mitochondrial and lysosomal volume, surface area, and mean signal intensity. Mitochondrial and lysosomal morphology were evaluated in a blinded manner across three independent biological replicates.

**In vivo protein and unfolded protein synthesis assay**

Protein synthesis rates were assessed in vivo using O-propargyl-puromycin (OP-Puro) incorporation as previously described. [5] Briefly, mice were administered OP-Puro at 49.5 mg/kg via intraperitoneal injection and sacrificed 1 hour post-injection. Bone marrow cells were harvested immediately following euthanasia and processed into single-cell suspensions. Cells were subsequently stained with surface antibodies as described above. Incorporated OP-Puro was detected using a Click-iT chemistry-based reaction according to the manufacturer's instructions (Thermo Fisher, Waltham, MA, USA), followed by flow cytometric analysis. Protein synthesis rates were quantified as fluorescence intensity within defined hematopoietic stem and progenitor cell populations. Data are normalized to internal wild-type control.

**Single-cell RNA sequencing (scRNA-seq)**

HSPCs were isolated by cell sorting from the bone marrow of adult offspring as described above. For each sample, cells were pooled from three mice derived from 3 independent litters (18,125 cells from Def, 24,953 cells from Control, 28,892 cells from Supp) and HSPCs were loaded onto the 10x Genomics Chromium Controller for single-cell capture and barcoding. Single-cell gene-expression libraries were prepared using the Chromium Next GEM Single Cell 3′ reagent kit (10x Genomics, Pleasanton, CA, USA) according to the manufacturer's instructions and sequenced on an Illumina NovaSeq platform to a target depth of approximately 200 million reads per sample. Raw base-call files were demultiplexed and converted to FASTQ format using Illumina bcl2fastq, and reads were aligned to the mouse reference genome (mm10) and quantified into gene-cell count matrices using 10x Genomics Cell Ranger. Count matrices from all samples were merged and imported into the R package Seurat (v4) for downstream analysis. Quality control was performed on a per-cell basis by computing the number of detected genes, total UMIs, and the proportion of mitochondrial reads; low-quality cells (fewer than ~500 detected genes or greater than ~10% mitochondrial reads) and putative doublets (cells with anomalously high gene and UMI counts) were removed.

Highly variable genes were identified and used for principal component analysis (PCA), and the top 30 principal components were used as input for Uniform Manifold Approximation and

Projection (UMAP) and shared-nearest-neighbor graph-based clustering. Cluster identities were assigned manually using previously described lineage and HSPC marker genes,[6,7] and the resulting annotated populations were used to resolve cell populations, relationships, and transcriptional changes across the folate dietary groups. Differential gene-expression analysis between conditions was performed using the Wilcoxon rank-sum test, and genes with an adjusted p-value <0.05 were considered significantly differentially expressed genes (DEGs). These gene sets were subjected to pathway-enrichment analysis querying the MSigDB Hallmark 2020, GO Biological Process 2023, and KEGG 2019 Mouse databases. In parallel, Gene Set Enrichment Analysis (GSEA; Broad Institute) was performed on genes pre-ranked by log2 fold-change against the MSigDB Hallmark and C2 curated gene-set collections, with gene sets at a false-discovery rate (FDR) <0.05 and a normalized enrichment score (NES) >1.2 considered significantly enriched. For visualization, marker- and signature-gene expression across clusters and folate dietary groups was summarized using bubble plots, in which dot size encodes the fraction of cells expressing each gene and color encodes scaled mean expression. DEGs were displayed as volcano plots of log2 FC versus −log10 adjusted p-value, and the per-cell expression of selected DEGs was projected across groups using violin plots. All visualization and statistical analyses were performed in Seurat using common R scripts (v4.4.1).

**Lactobacillus casei assay to quantify total folate concentrations:**

Total folate concentrations in plasma and whole tissues were quantified by *Lactobacillus casei* microbiological assay as previously described,[8] with bacterial growth quantified at 600 nm on Epoch Microplate Spectrophotometer (Biotek Instruments). Tissue folate concentrations were normalized to protein concentration, as assessed by Lowry-Bensadoun assay. [9]

**Statistical analysis**

All statistical analyses were performed using GraphPad Prism (GraphPad Software). Data are presented as mean ± standard deviation (SD) unless otherwise indicated. Biological replicates (n) are defined as independent samples derived from distinct litters, unless otherwise specified. Technical replicates are indicated in the corresponding figure legends where applicable.

For comparisons between two groups, unpaired two-tailed Student's t-tests were used. For comparisons involving more than two groups, one-way analysis of variance (ANOVA) followed by multiple comparisons testing against the control group (Dunnett's test) was performed. A p-value of < 0.05 was considered statistically significant. Data distribution was assumed to be normal but was not formally tested. No data points were excluded.

**ATAC Seq**

ATACseq data were processed using the nf-core/atacseq pipeline (v 2.1.2),[10] which follows best practices for chromatin accessibility data analysis. The pipeline performs adaptor trimming (Trim Galore!), read alignment to the mm10 genome (Bowtie2), duplicate marking (picard), duplicate removal (SAMtools), peak calling (MACS2), and quality control metrics (ataqv). One replicate (hsc_0mg_REP1) showed lower FRiP and peak counts relative to other samples, likely reflecting reduced library complexity due to low cell input and sample handling constraints inherent to primary HSCs. This replicate was retained in the analysis but flagged; results were confirmed to be consistent with the remaining replicates at the consensus peak level. DESeq2 was used to assess differential accessibility from the consensus peaks in all conditions (log2 fold change >1; p-value <0.05).[11] Individual regions of interests were manually visualized with IGV for replicate consistency and signal to background enrichment. Known transcription factor motif enrichment was performed on differentially accessible regions (DARs) using Homer (v5.1; `findMotifsGenome.pl -size given`) against the mm10 genome. Signal heatmaps over DARs were generated using deepTools (v3.5.6). Size factor-normalized bigwig files were used as input. For each pairwise comparison, `computeMatrix reference-point` was run centering on the peak midpoint (±2kb), with missing data set to zero and all-zero regions skipped. Heatmaps were rendered with `plotHeatmap`. Additional data visualization was performed with R (v4.4.1).

**Data Availability**

Datasets generated during this study are available at Gene Expression Omnibus under ascension numbers GSE341612 and GSE341655

#### Supplementary Tables

**Table S1. Composition of control, deficient and supplemented diets.** Diets were obtained from Dyets Inc., Bethlehem PA.

| Dyet #117814GI Custom 2 mg/kg Folic Acid AIN-93G Based Diet | | | |
|---|---|---|---|
| **Ingredient** | **kcal/gram** | **grams/kg** | **kcal/kg** |
| Sterile Casein | 3.58 | 200 | 716 |
| L-Cystine | 4 | 3 | 12 |
| Sucrose | 4 | 100 | 400 |
| Cornstarch | 3.6 | 395.5 | 1423.7 |
| Dyetrose | 3.8 | 132 | 501.6 |
| Soybean Oil | 9 | 70 | 630 |
| t-Butylhydroquinone | 0 | 0.014 | 0 |
| Cellulose | 0 | 50 | 0 |
| Mineral Mix #210025 | 0.88 | 35 | 30.8 |
| Folic Acid Premix (1mg/kg Folate) | 4 | 2 | 8 |
| Choline Bitartrate | 0 | 2.5 | 0 |
| Vitamin Mix #317761 (no Folate) | 3.87 | 10 | 38.7 |

| Dyet #117814GI Custom 0 mg/kg Folic Acid AIN-93G Based Diet | | | |
|---|---|---|---|
| **Ingredient** | **kcal/gram** | **grams/kg** | **kcal/kg** |
| Sterile Casein | 3.58 | 200 | 716 |
| L-Cystine | 4 | 3 | 12 |
| Sucrose | 4 | 100 | 400 |
| Cornstarch | 3.6 | 395.5 | 1423.7 |
| Dyetrose | 3.8 | 132 | 501.6 |
| Soybean Oil | 9 | 70 | 630 |
| t-Butylhydroquinone | 0 | 0.014 | 0 |
| Cellulose | 0 | 50 | 0 |
| Mineral Mix #210025 | 0.88 | 35 | 30.8 |
| Folic Acid Premix (1mg/kg Folate) | 0 | 0 | 0 |
| Choline Bitartrate | 0 | 2.5 | 0 |
| Vitamin Mix #317761 (no Folate) | 3.87 | 10 | 38.7 |

| Dyet #117814GI Custom 8 mg/kg Folic Acid AIN-93G Based Diet | | | |
|---|---|---|---|
| **Ingredient** | **kcal/gram** | **grams/kg** | **kcal/kg** |
| Sterile Casein | 3.58 | 200 | 716 |
| L-Cystine | 4 | 3 | 12 |
| Sucrose | 4 | 100 | 400 |
| Cornstarch | 3.6 | 395.5 | 1423.7 |
| Dyetrose | 3.8 | 132 | 501.6 |
| Soybean Oil | 9 | 70 | 630 |
| t-Butylhydroquinone | 0 | 0.014 | 0 |
| Cellulose | 0 | 50 | 0 |
| Mineral Mix #210025 | 0.88 | 35 | 30.8 |
| Folic Acid Premix (1mg/kg Folate) | 16 | 8 | 32 |
| Choline Bitartrate | 0 | 2.5 | 0 |
| Vitamin Mix #317761 (no Folate) | 3.87 | 10 | 38.7 |

**Table S2. Antibodies used in this study**

| Antibody | Source | Identifier |
|---|---|---|
| CD45 BUV395 | BD Biosciences | RRID:AB_2916597 (BD Biosciences Cat. No. 564176) |
| C-Kit APC-Cy7 | Biolegend | RRID AB_1626278 (BioLegend Cat. No. 105826) |
| Sca1 Pacific Blue | Biolegend | RRID AB_2143237 (BioLegend Cat. No. 122520) |
| Flk2 APC | Biolegend | RRID AB_2107050 (BioLegend Cat. No. 135310) |
| CD48 BV605 | Biolegend | RRID AB_2650825 (BioLegend Cat. No. 103441) |
| CD150 BV785 | Biolegend | RRID AB_2565962 (BioLegend Cat. No. 115937) |
| IL7Ra PE-Cy7 | Biolegend | RRID AB_2621859 (Tonbo Biosciences Cat. No. 60-1271) |
| CD3 Biotin | Biolegend | RRID AB_312669 (BioLegend Cat. No. 100304) |
| CD4 Biotin | Biolegend | RRID AB_312689 (BioLegend Cat. No. 100404) |
| CD5 Biotin | eBioscience | RRID AB_466340(Thermo Fisher Scientific Cat. No. 13-0051-85) |
| CD8 Biotin | Biolegend | RRID AB_312743 (BioLegend Cat. No. 100704) |
| CD19 Biotin | Biolegend | RRID AB_313639 (BioLegend Cat. No. 115504) |
| NK1.1 Biotin | Biolegend | RRID AB_313391 (BioLegend Cat. No. 108704) |
| Ter119 Biotin | Biolegend | RRID AB_313705 (BioLegend Cat. No. 116204) |
| FceRIa Biotin | Biolegend | RRID:AB_1626106 (BioLegend Cat. No. 134304) |
| F4/80 Biotin | Biolegend | RRID AB_893501 (BioLegend Cat. No. 123106) |
| Gr1 Biotin | Tonbo Biosciences | RRID AB_2621652 (Tonbo Biosciences Cat. No. 30-5931) |
| CD45.1 BV605 | Biolegend | RRID:AB_2562565 (BioLegend Cat. No. 110738) |
| CD45.2 BV785 | Biolegend | RRID AB_2562604 (BioLegend Cat. No. 109839) |
| Ter119 PE-Cy5 | eBioscience | RRID AB_468811 (Thermo Fisher Scientific Cat. No. 15-5921-83) |
| B220 APC-Cy7 | Biolegend | RRID:AB_313007 |

| | | |
|---|---|---|
| | | (BioLegend Cat. No. 103224) |
| CD3 APC | Biolegend | RRID AB_312677 (BioLegend Cat. No. 100312) |
| CD11b PE-Cy7 | Tonbo Biosciences | RRID AB_2621836 (Tonbo Biosciences Cat. No. 60-0112) |
| Gr1 v450 | Tonbo Biosciences | RRID AB 2621968 (Tonbo Biosciences Cat. No. 75-5931) |
| CD61 PE-Cy7 | Biolegend | RRID:AB_2687361 (BioLegend Cat. No. 104318) |
| CD105 PE-Cy7 | eBioscience | RRID AB_2573380 (Thermo Fisher Scientific Cat. No. 25-1051-82) |
| CD41 BV421 | Biolegend | RRID AB_2650893 (BioLegend Cat. No. 133912) |
| CD16/32 APC | Biolegend | RRID AB_1953273 (BioLegend Cat. No. 101326) |
| Streptavadin PE-Cy5 | Biolegend | Cat. No. 405205 |
| Streptavadin BUV395 | BD Biosciences | RRID:AB_2869553 (BD Biosciences Cat. No. 564176) |
| CD150 PE-Cy7 | Biolegend | RRID AB_756199 (BioLegend Cat. No. 122514) |
| CD3 FITC | BioLegend | Cat #100204; RRID:AB_312661 |
| CD8A FITC | eBioscience | Cat #11-0081-85; RRID:AB_464916 |
| CD5 FITC | BioLegend | Cat #100606; RRID:AB_312735 |
| GR1 FITC | BioLegend | Cat #108406; RRID:AB_313371 |
| TER119 FITC | Biolegend | Cat #116206; RRID:AB_313709 |
| B220 FITC | eBioscience | Cat #11-0452-85; RRID:AB_465055 |
| CD48 APC | Biolegend | RRID AB_571997 (BioLegend Cat. No. 103412) |
| CKIT APC-EFLUOR 780 | eBioscience | Cat #47-1171-82; RRID:AB_1272177 |
| SCA1 PERCPCY5.5 | eBioscience | Cat #45-5981-82; RRID:AB_914372 |
| CD150 PE | BioLegend | Cat #115904; RRID:AB_313683 |
| CD4 FITC | BioLegend | Cat #100406; RRID:AB_312691 |
| CD11B FITC | eBioscience | Cat #12-0112-83; RRID:AB_2734870 |
| CD48 FITC | BioLegend | Cat #103404; RRID:AB_313019 |

| | | |
|---|---|---|
| CD34 ALEXAFLUOR700 | eBioscience | Cat #56-0341-82; RRID:AB_493998 |
| CD16/32 PERCPCY5.5 | BioLegend | Cat #101324; RRID:AB_1877267 |
| CD11B APC | eBioscience | Cat #47-0112-82; RRID:AB_1603093 |
| CD11b-Biotin | Biolegend | RRID AB_312787 (BioLegend Cat. No. 101204) |
| Anti-CDC42-GTP Monoclonal Antibody | NewEast Biosciences | Cat# 26905, RRID:AB_1961759 |
| Recombinant Anti-LAMP1 antibody [EPR21026] | Abcam | Cat# ab208943, RRID:AB_2923327 |
| Anti-TOMM20 antibody [EPR15581-39] | Abcam | Cat# ab283317, RRID:AB_3665348 |
| Recombinant Anti-DRP1 antibody [EPR19274] | Abcam | Cat# ab219596, RRID:AB_3065099 |
| Hoechst 33342 | Abcam | Cat# ab228551, RRID:AB_3720789 |
| TPE-MI | MedChem Express | Cat# HY-143218 |
| DCFDA | Abcam | Cat# ab113851 |
| BODIPY | ThermoFisher | Cat# D3922 |
| TMRM | Thermo Scientific | Cat# M20036 |

**Table S3. Chemicals, Peptides and Reagents used in this study**

| Resource | Source | Identifier |
|---|---|---|
| ACK Lysing Buffer | Thermofisher | Thermofisher Scientific Cat. No. A1049201 |
| Calibrite Beads - APC | BD Biosciences | BD Biosciences Cat. No. 340487 |
| CD117 Microbeads, Mouse | Miltenyi Biotech | Miltenyi Biotech Cat. No. 130-091-224 |
| DAPI | BD Biosciences | BD Biosciences Cat. No. 564907 |
| DMSO | Millipore Sigma | Millipore Sigma Cat. No D2650 |
| DPBS (1X), no calcium, no magnesium | Fisher | Fisher Cat. No. 14190250 |
| EDTA | Fisher | Fisher Cat. No. BP1201 |
| FBS | Gibco | Thermo Fisher Scientific Cat. No. 26140079 |
| Hoescht 33342 | Invitrogen | Thermo Fisher Scientific Cat. No. H3570 |
| Propidium Iodide | VWR | WVR Cat. No. 89139-064 |
| StemPro-34 (1X) | Thermo Fisher | 10639011 |
| L-Glutamine | Thermo Fisher | 25030149 |
| Primocin | Fisher | NC9141851 |
| SCF | Peprotech | 250-03 |
| Flt3L | Peprotech | 250-31L |
| TPO | Peprotech | 315-14 |
| Alexa Fluor 555 azide | Thermo Fisher Scientific | cat. no. A20012 |
| O-propargyl-puromycin | OP-Puro; Medchem Source | |
| ProLong™ Diamond Antifade Mountant | Invitrogen | P36965 |
| Beta-mercaptoethanol | Sigma | M6250 |
| Folic acid Casei Medium | HIMEDIA | M543 |
| HEPES | VWR | BP310 |
| CHES | Alfa Aesar | A18047 |
| Rat serum | VWR | 103219-662 |
| Sodium ascorbate | VWR | A7631 |

**Table S4. Experimental models used in this study**

| Resource | Source | Identifier |
|---|---|---|
| Mouse: C57BL/6J | The Jackson Laboratory | RRID:IMSR_JAX:006965 |
| Mouse: mT/mG | The Jackson Laboratory | RRID:IMSR_JAX:007576 |
| Mouse: Flk2-Cre (Flt3-Cre) | N/A | T. Boehm and C. Bleul (Max Planck, Freiburg, Germany) |
| Mouse: B6 CD45.1 | The Jackson Laboratory | RRID:IMSR_JAX:002014 |
| Lacticaseibacillus rhamnosus | ATCC | 7469 |

**Table S5. Kits used in this study**

| Resource | Source | Identifier |
|---|---|---|
| Click-iT Cell Reaction Buffer Kit | Thermo Fisher Scientific | cat. no. C10269 |

**Table S6. Software used in this study**

| | | |
|---|---|---|
| Flowjo v.10 | Treestar | RRID:SCR_000410 |
| Graphpad Prism v.10 | Graphpad | https://www.graphpad.com |
| ImageJ | NIH ImageJ | RRID:SCR_003070 |
| Imaris | Oxford Instruments | RRID:SCR_007370 |
| Zen Lite | ZEISS | RRID:SCR_023747 |

#### Supplemental Figures

**Figure S1. Folate levels and erythropoiesis in response to maternal folate nutrition.**

**A)** Maternal blood plasma folate concentration at 14.5 days post-coitum (dpc) measured by microbiological assay.

**B)** Folate concentration (normalized per ug of protein) in the fetal liver at embryonic day (E)14.5 microbiological assay.

**C-F)** Maternal blood red blood cell quantification (million per μl) (**C**), hemoglobin (grams per deciliter) (**D**) hematocrit (percent of whole blood) (**E**), and white blood cell quantification (thousand per μl) (**F**).

**G)** Red blood cell quantification in the fetal liver at E14.5 (million per μl).

**H-J)** Red blood cell quantification (million per μl) (**H**), hemoglobin (grams per deciliter) (**I**), and hematocrit (percent of whole blood) (**J**) at postnatal day 14.

**K-M)** Red blood cell quantification (million per μl) (**K**), hemoglobin (grams per deciliter) (**L**) and hematocrit (percent of whole blood) (**M**) in adult offspring.

**N)** Intracellular folate concentrations in long-term hematopoietic stem cells (HSCs) at embryonic day 14.5 measured by mass spectrometry-based metabolomic profiling.

**O)** Intracellular folate concentrations in HSCs isolated from adult offspring measured by mass spectrometry-based metabolomic profiling.

N = 3 independent experiments representing three independent litters per group. Significance determined by one-way ANOVA with multiple comparisons or unpaired student's t-test for; *P < 0.05; **P < 0.01.

**Figure S2. Hematopoietic outcomes of prenatal folate status in situ and upon transplantation.**

**A-E)** Total cellularity of adult offspring bone marrow hematopoietic cell populations quantified by flow cytometry. (**A**) common lymphoid progenitors (CLPs), (**B**) B-cells, (**C**) T-cells, (**D**) megakaryocyte progenitors (MkPs), (**E**) Granulocytes.

**F-H)** Total cellularity of adult offspring peripheral blood hematopoietic cell populations quantified by flow cytometry. (**F**) granulocyte/macrophages (GMs), (**G**), B-cells and (**H**) T-cells.

**I)** Bone marrow chimerism in LT-HSCs, ST-HSCs, $MPP4^{Ly}$, $MPP3^{GM}$ and $MPP2^{MkE}$ in primary recipients of 200 HSCs isolated from Def adult offspring bone marrow. Gold data points indicate primary transplantation recipients used as donors for secondary transplantation in Fig 1K.

**J)** Bone marrow chimerism in LT-HSCs, ST-HSCs, $MPP4^{Ly}$, $MPP3^{GM}$ and $MPP2^{MkE}$ in primary recipients of 200 HSCs isolated from Supp adult offspring bone marrow. Gold data points indicate primary transplantation recipients used as donors for secondary transplantation in Fig 1L.

N = 3 independent experiments representing three independent litters per group. Significance determined by one-way ANOVA with multiple comparisons or unpaired student's t-test for analyses. *P < 0.05; **P < 0.01; ***P<0.001; ****, P<0.0001.

**Figure S3. Overview of changes to the HSC metabolic landscape.**

**A)** Principal component analysis (PCA) scores of metabolite distribution in adult offspring HSCs analyzed by MetaboAnalyst.

**B)** Heatmap of the top 50 metabolites by sample variability in adult offspring HSCs analyzed by MetaboAnalyst.

**C)** Volcano plot of all differentially expressed metabolites in folate Def adult offspring HSCs compared to control.

**D)** Volcano plot of all differentially expressed metabolites in folate Supp adult offspring HSCs compared to Control.

**E)** Representative electron microscopy images of Def, Control and Supp HSCs, taken at 15,000x magnification.

N = 3 independent experiments representing three independent litters per group.

**Figure S4. Unsupervised clustering identified discrete populations corresponding to long-term HSCs, multipotent progenitors, and lineage-primed progenitors.**

**A)** Nineteen discrete populations identified within sorted $Lin^{-}Sca1^{+}cKit^{+}$ (LSK) bone marrow cells identified by unsupervised clustering of single-cell RNA sequencing (scRNA-Seq).

**B)** Frequency of each scRNAseq cluster in Folate Def, Folate Supp, and Control offspring LSK cells.

N = 3 independent experiments representing three independent litters per group.

**Figure S5. Imaris-based 3D reconstruction and quantification of lysosomes and mitochondrial-associated proteins.**

**A)** Representative workflow for 3D reconstruction. A 2D confocal slice showing DAPI (blue), TOM20 (red), and LAMP1 (green) staining is used to generate 3D renderings of mitochondrial (TOM20) and lysosomal (LAMP1) compartments. Intracellular TOM20 signal is used to define the cell boundary (blue), enabling reconstruction of the cell surface and spatial mapping of LAMP1$^{+}$ lysosomes within the cellular volume.
**B)** Quantification of total mitochondrial signal normalized to cell size. TOM20 fluorescence intensity was measured within the reconstructed cellular volume and normalized to either cell surface area or total cell volume to account for differences in cell size.
**C)** 3D reconstruction of TOM20 (red) and CDC42 (green). Representative confocal images (DAPI, TOM20, CDC42) were rendered in 3D. The intracellular TOM20 signal was used to define the cell boundary, enabling spatial localization of CDC42 within the reconstructed cellular volume.
N = 3 independent experiments representing three independent litters per group. Significance determined by unpaired student’s t-test.

**Figure S6. Metabolic alterations in fetal liver hematopoiesis in response to prenatal folate.**

**A-E)** Fetal liver hematopoietic cell populations quantified by flow cytometry: (**A**) MPP2$^{MkE}$ (**B**) MPP3$^{GM}$, (**C**) MPP4$^{Ly}$, (**D**) CD11b$^{+}$, (**E**) B-cells.

**F)** Seahorse extracellular acidification rate (ECAR) performed on fetal liver HSPCs, normalized to 1000 cells.

**G)** Seahorse oxygen consumption rate (OCR), performed on fetal liver HSPCs, normalized to 1000 cells.

**H-J)** Mean fluorescence intensity (MFI) of tetramethylrhodamine (TMRM) staining in fetal liver (**H**) LT-HSC, (**I**) ST-HSC, (**J**) MPP.

N = 3 independent experiments representing three independent litters per group. Significance determined by one-way ANOVA with multiple comparisons; * $P < 0.05$; ** $P < 0.01$; **** $P<0.0001$.

**Figure S7. Metabolomic profiling of fetal liver HSCs in response to varying prenatal folate**

**A)** Heatmap of the top 50 metabolites by sample variability in fetal liver HSCs analyzed by MetaboAnalyst.
**B**) Normalized relative abundances of select individual metabolites representing glucose metabolism from untargeted metabolomics analysis: Glucose, Glutamine, Glutamate.

**C**) Normalized relative abundances of select individual metabolites representing methionine metabolism from untargeted metabolomics analysis: Methionine, Methionine S-oxide.

**D)** Normalized relative abundances of selected individual metabolites representing long-chain fatty acid metabolism from untargeted metabolomics analysis: lauric acid, stearic acid, oleic acid, linoleic acid, eicosatetraenoic acid.

N = 3 independent experiments representing three independent litters per group. Significance determined by unpaired student's t-test. *P < 0.05; **P < 0.01.

**Figure S8. Supplementary chromatin accessibility analyses in adult HSCs**

**A)** Principal component analysis (PCA) scores of differentially accessible region (DAR) distribution in adult offspring HSCs.
**B-C)** Transcription start site (TSS) enrichment profiles for ATAC-seq libraries from Def (**B**) and Supp (**C**) adult offspring HSCs.

**D-E)** Genomic distribution of DARs identified in Def (**D**) and Supp (**E**) adult offspring HSCs. Stacked bar plots indicate the proportion of DARs associated with gene regions.

**F-G)** Enriched transcription factor (TF) binding motifs in closed DARs in Def vs Control (**F**) and Supp vs Control (**G**) adult offspring HSCs.

**H)** Genes with NRF2/NF-E2 motif-containing DARs in Def vs Control adult offspring HSCs.
**I)** DARs of Polycomb-associated genes in Supp vs Control adult offspring HSCs.
**J)** DARs of self-renewal associated genes in Supp vs Control adult offspring HSCs.
**K)** DARs of mitochondrial pyruvate handling associated genes in Supp vs Control adult offspring HSCs.
N = 3 independent experiments representing three independent litters per group

**Supplemental References**


1. Benz, C., Martins, V.C., Radtke, F., and Bleul, C.C. (2008). The stream of precursors that colonizes the thymus proceeds selectively through the early T lineage precursor stage of T cell development. J Exp Med *205*, 1187-1199. 10.1084/jem.20072168.
2. Muzumdar, M.D., Tasic, B., Miyamichi, K., Li, L., and Luo, L. (2007). A global double-fluorescent Cre reporter mouse. Genesis *45*, 593-605. 10.1002/dvg.20335.
3. Nemkov, T., Reisz, J.A., Gehrke, S., Hansen, K.C., and D'Alessandro, A. (2019). High-Throughput Metabolomics: Isocratic and Gradient Mass Spectrometry-Based Methods. Methods Mol Biol *1978*, 13-26. 10.1007/978-1-4939-9236-2_2.
4. Hinge, A., He, J., Bartram, J., Javier, J., Xu, J., Fjellman, E., Sesaki, H., Li, T., Yu, J., Wunderlich, M., et al. (2020). Asymmetrically Segregated Mitochondria Provide Cellular Memory of Hematopoietic Stem Cell Replicative History and Drive HSC Attrition. Cell Stem Cell *26*, 420-430.e426. 10.1016/j.stem.2020.01.016.
5. Hidalgo San Jose, L., and Signer, R.A.J. (2019). Cell-type-specific quantification of protein synthesis in vivo. Nat Protoc *14*, 441-460. 10.1038/s41596-018-0100-z.
6. Kucinski, I., Campos, J., Barile, M., Severi, F., Bohin, N., Moreira, P.N., Allen, L., Lawson, H., Haltalli, M.L.R., Kinston, S.J., et al. (2024). A time- and single-cell-resolved model of murine bone marrow hematopoiesis. Cell Stem Cell *31*, 244-259.e210. 10.1016/j.stem.2023.12.001.
7. Cabezas-Wallscheid, N., Klimmeck, D., Hansson, J., Lipka, D.B., Reyes, A., Wang, Q., Weichenhan, D., Lier, A., von Paleske, L., Renders, S., et al. (2014). Identification of regulatory networks in HSCs and their immediate progeny via integrated proteome, transcriptome, and DNA methylome analysis. Cell Stem Cell *15*, 507-522. 10.1016/j.stem.2014.07.005.
8. Suh, J.R., Oppenheim, E.W., Girgis, S., and Stover, P.J. (2000). Purification and properties of a folate-catabolizing enzyme. J Biol Chem *275*, 35646-35655. 10.1074/jbc.M005864200.
9. Bensadoun, A., and Weinstein, D. (1976). Assay of proteins in the presence of interfering materials. Anal Biochem *70*, 241-250. 10.1016/s0003-2697(76)80064-4.
10. Ewels, P.A., Peltzer, A., Fillinger, S., Patel, H., Alneberg, J., Wilm, A., Garcia, M.U., Di Tommaso, P., and Nahnsen, S. (2020). The nf-core framework for community-curated bioinformatics pipelines. Nat Biotechnol *38*, 276-278. 10.1038/s41587-020-0439-x.
11. Love, M.I., Huber, W., and Anders, S. (2014). Moderated estimation of fold change and dispersion for RNA-seq data with DESeq2. Genome Biol *15*, 550. 10.1186/s13059-014-0550-8.

**Supplement Figure 1 – Folate levels and erythropoiesis in response to maternal folate nutrition**

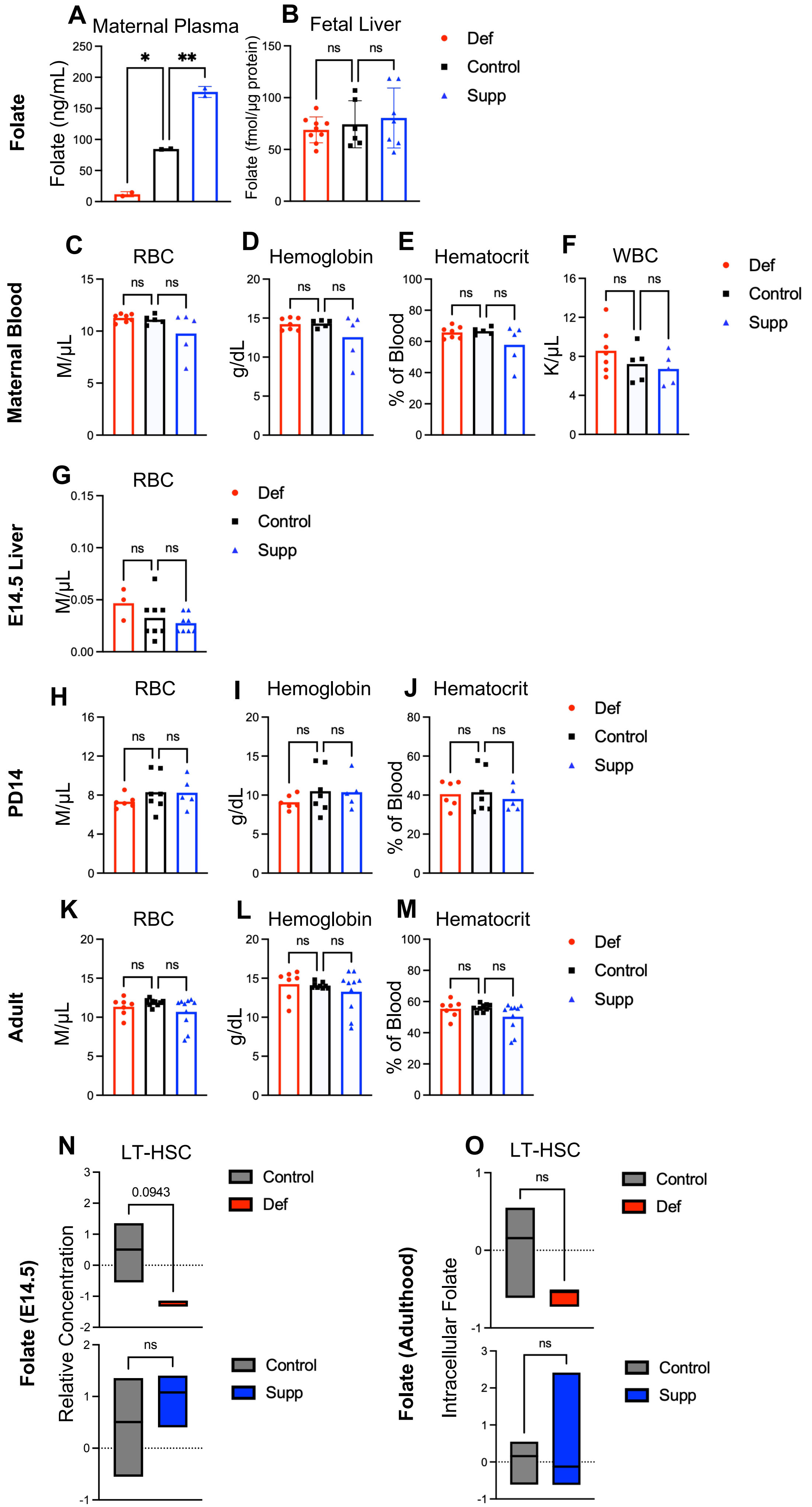

**Supplementary Figure 2 – Hematopoietic outcomes of prenatal folate status in situ and upon transplantation**

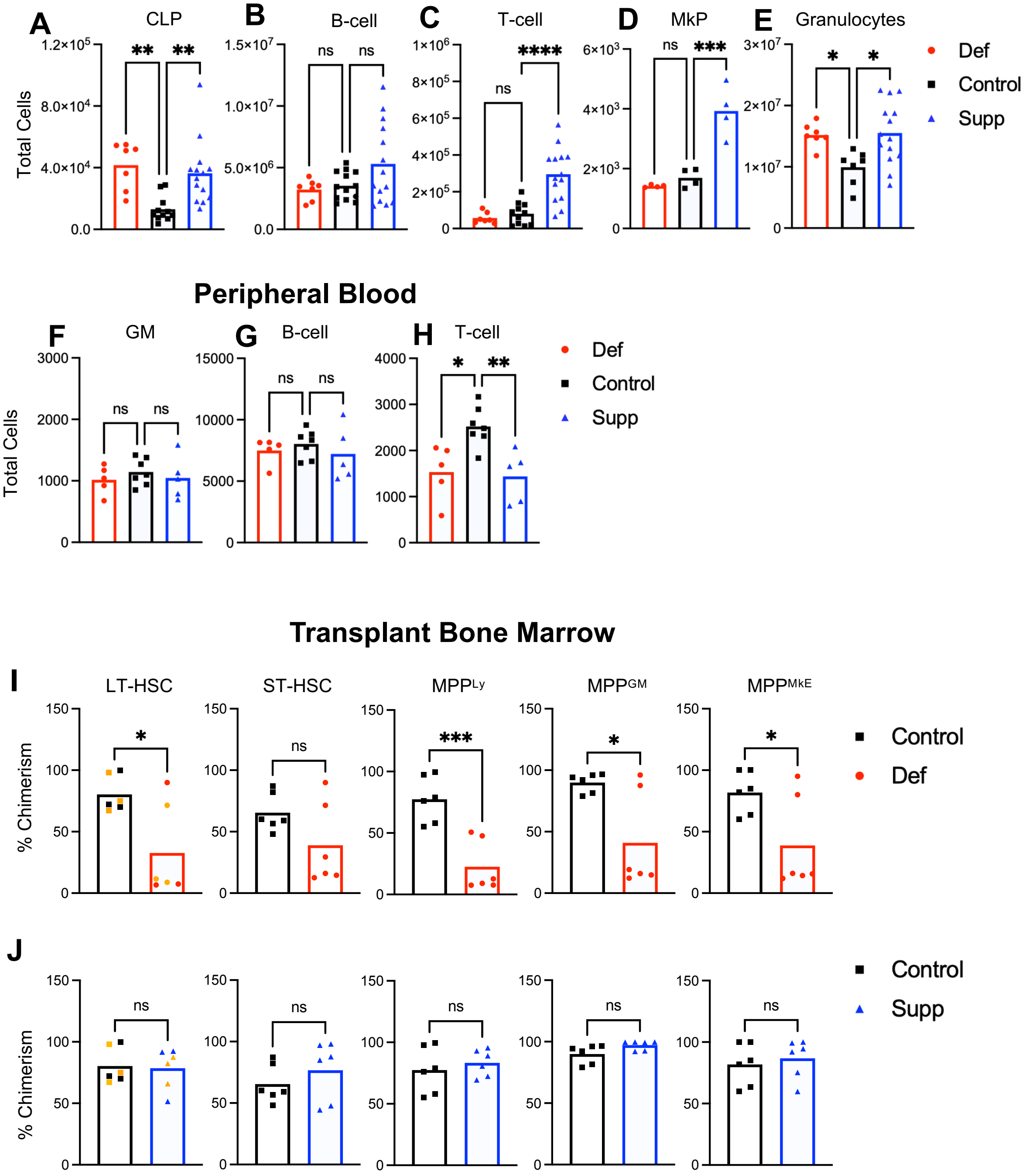

### Supplementary Figure 3 – Overview of changes to the HSC metabolic landscape

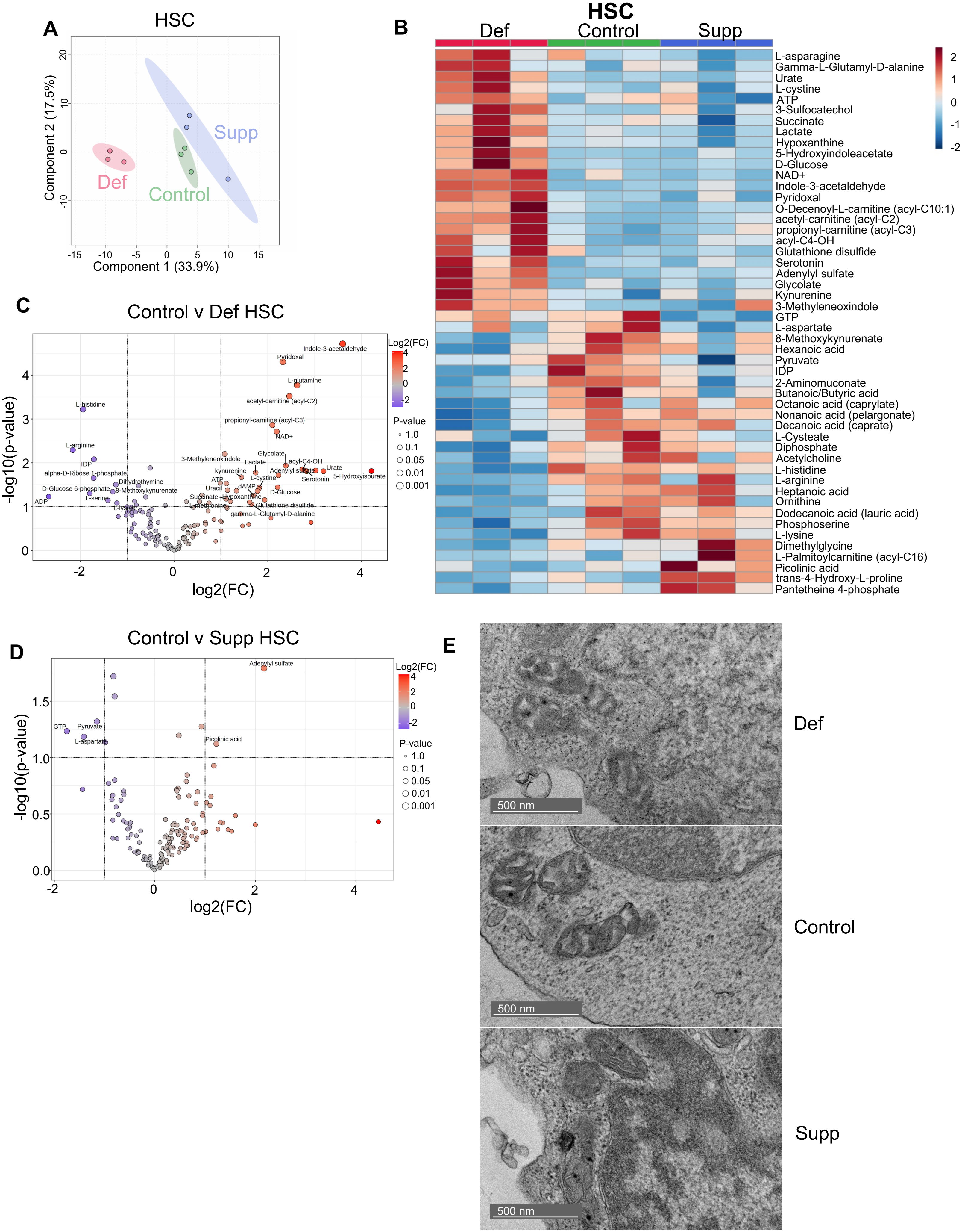

**Supplementary Figure 4 – Unsupervised clustering identified discrete populations corresponding to long-term HSCs, multipotent progenitors, and lineage-primed progenitors**

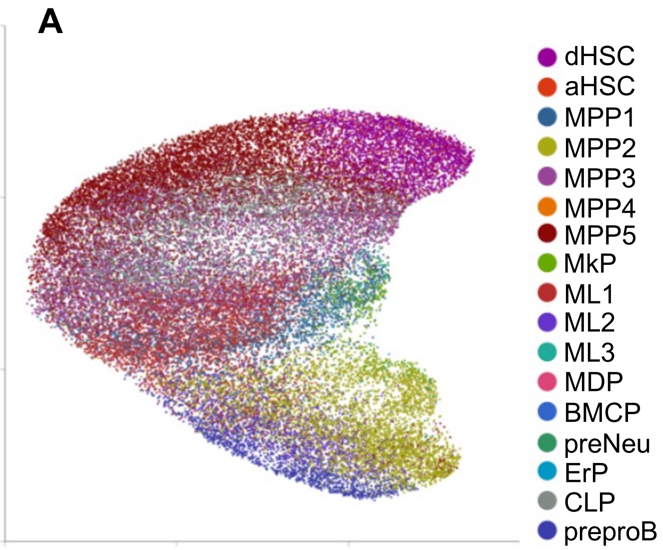


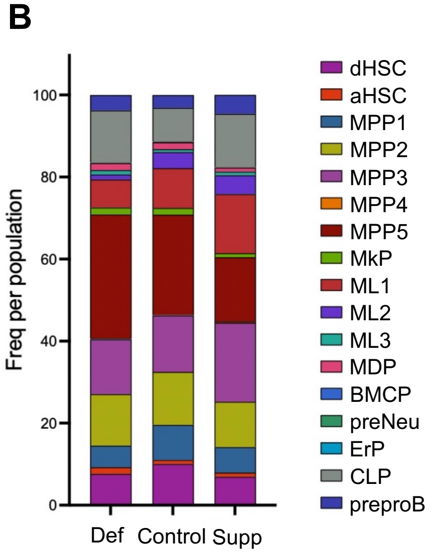

**Supplementary Figure 5 – Imaris-based 3D Reconstruction and Quantification of Lysosomes and Mitochondrial-Associated Proteins**

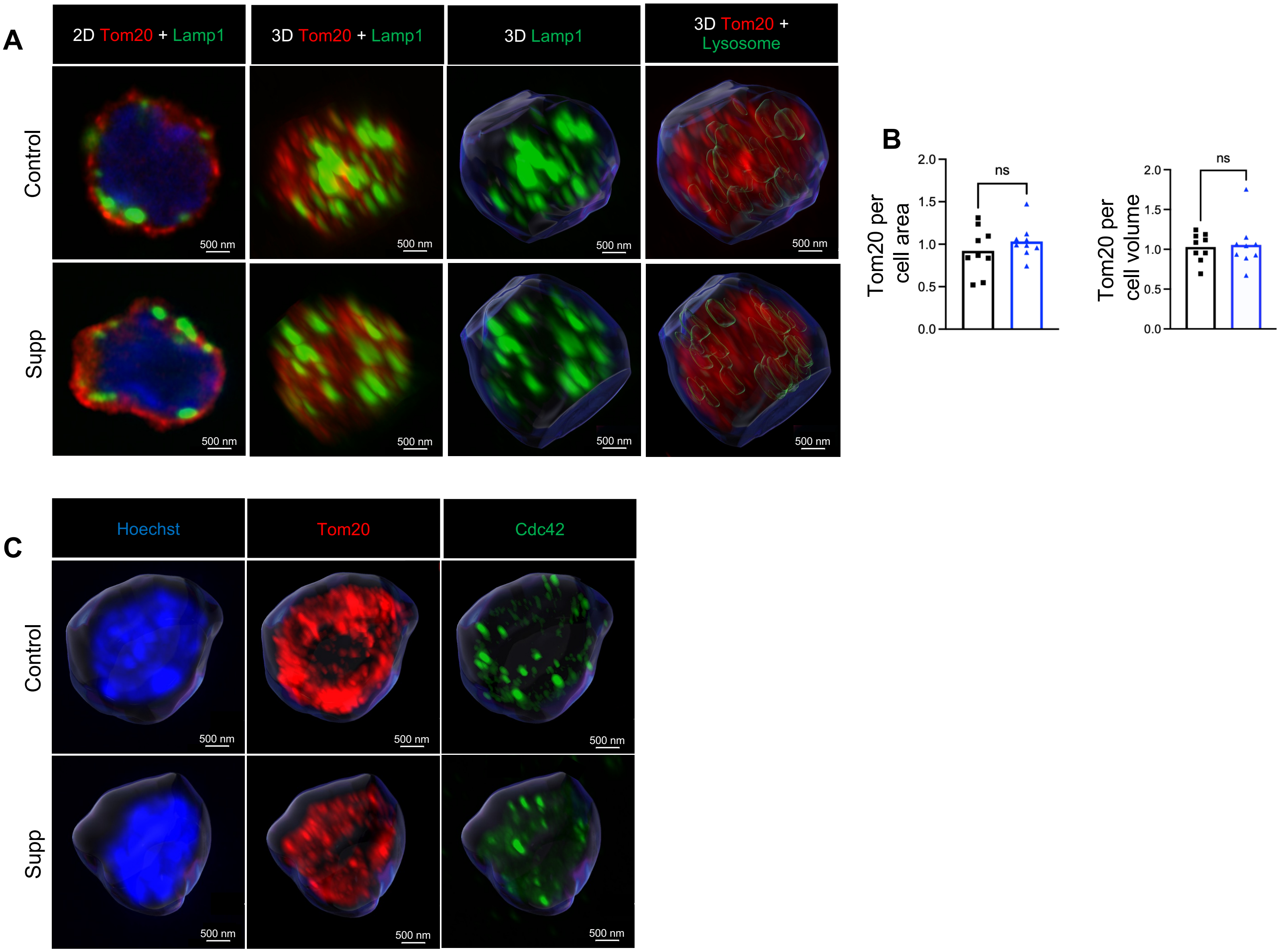

**Supplement Figure 6 – Metabolic alterations in fetal liver hematopoiesis in response to prenatal folate**

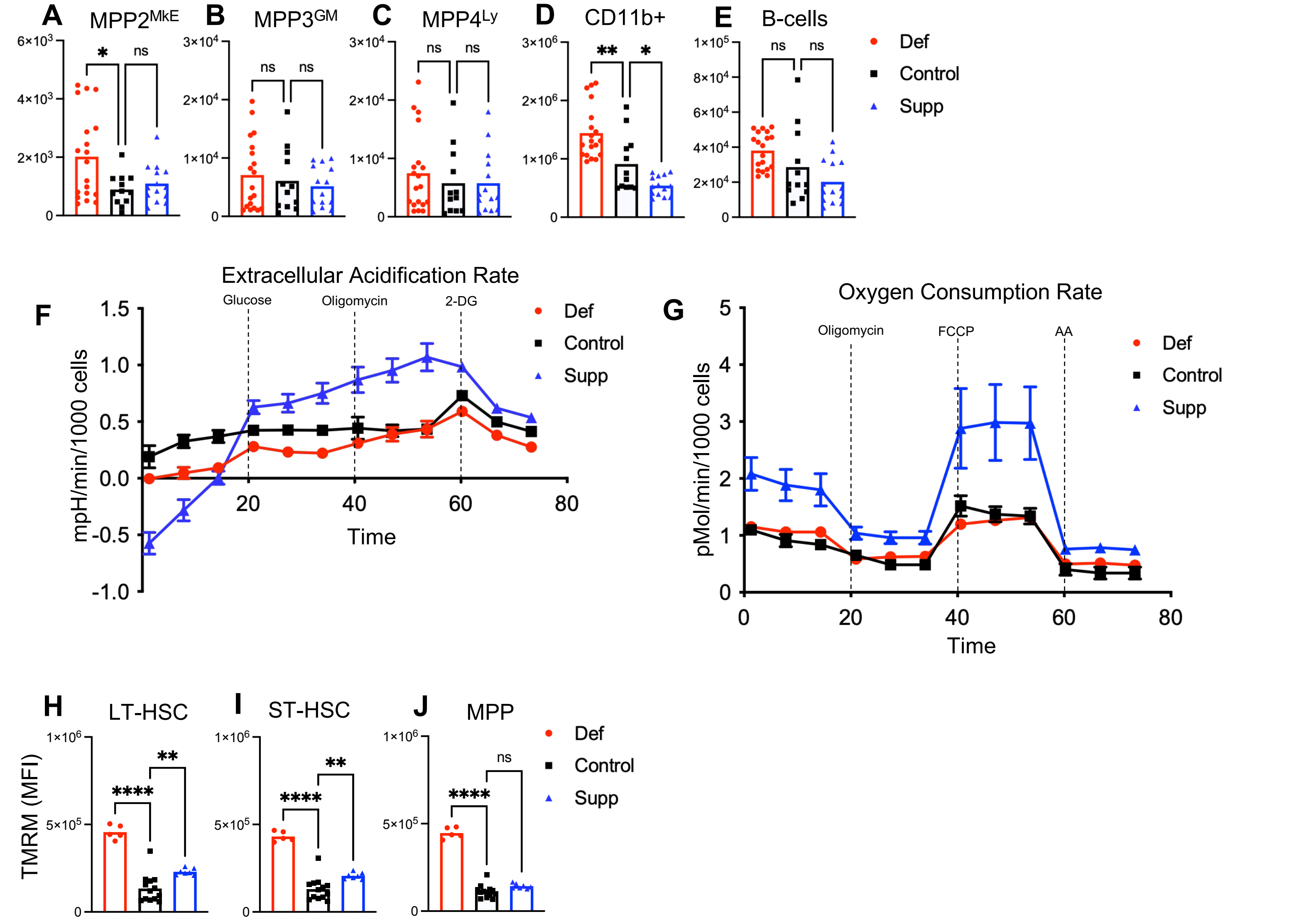

**Supplement Figure 7 – Metabolomic profiling of fetal liver HSCs in response to varying prenatal folate**

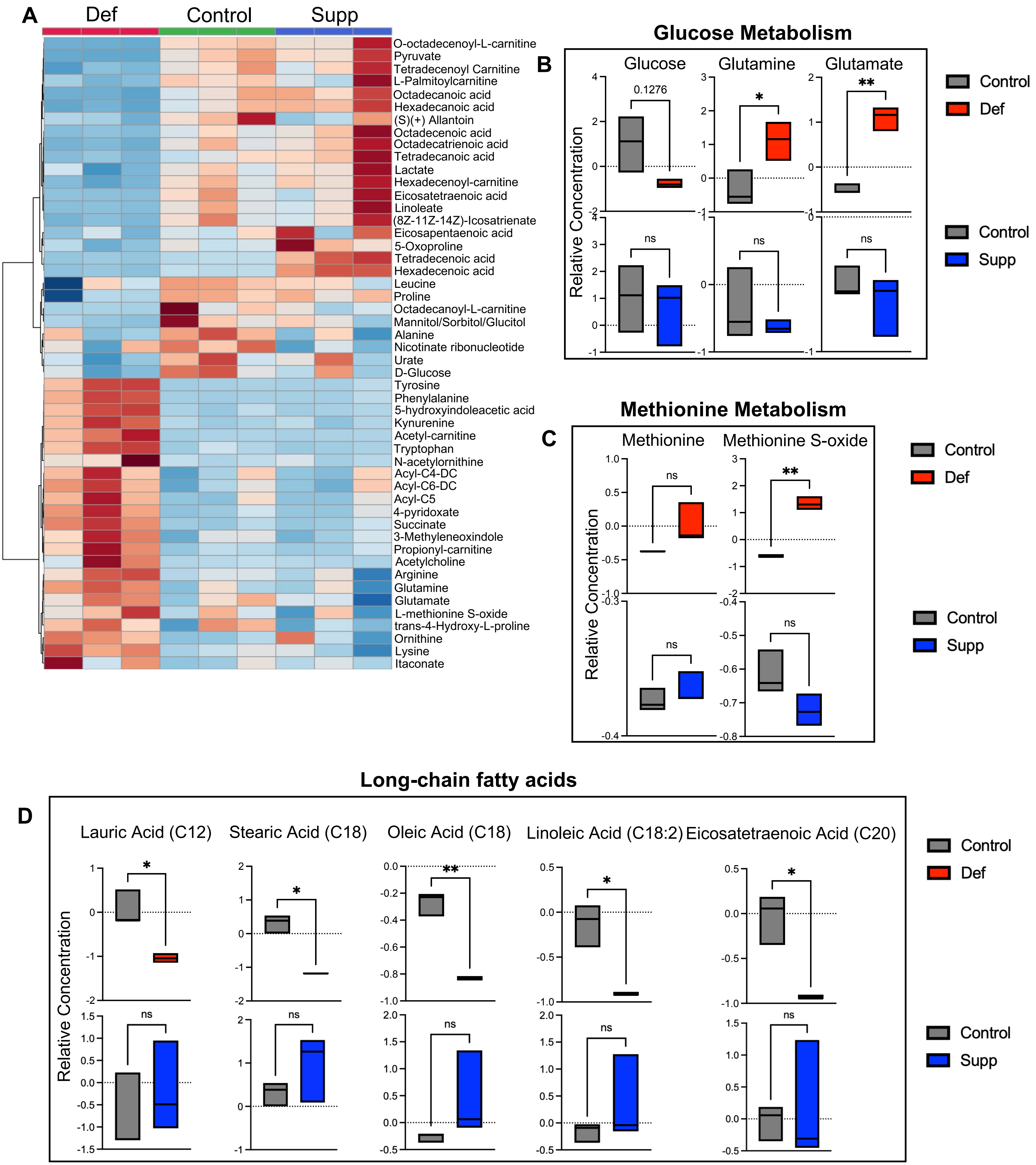

### Supplement Figure 8 – Supplementary chromatin accessibility analyses in adult HSCs

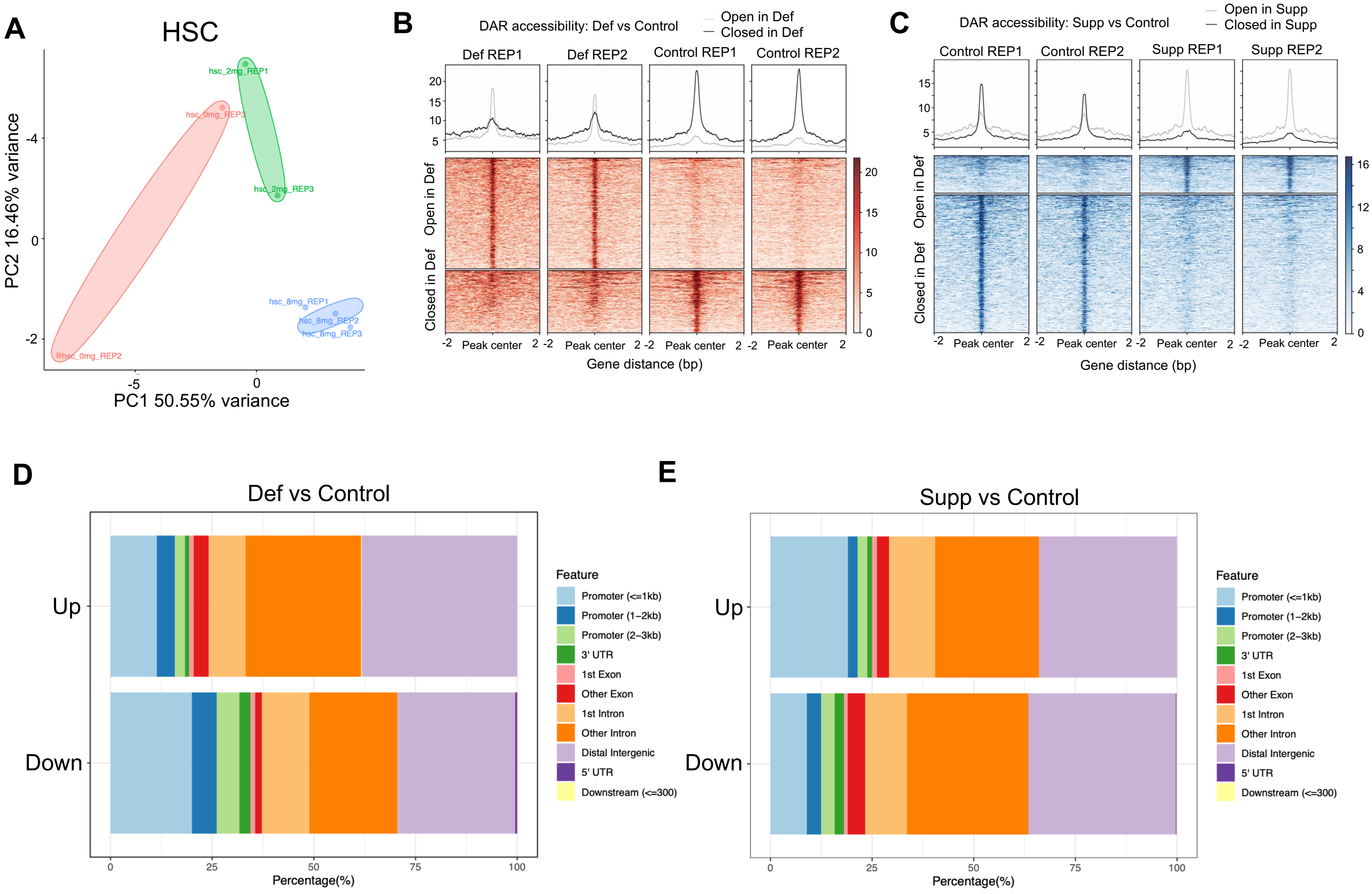


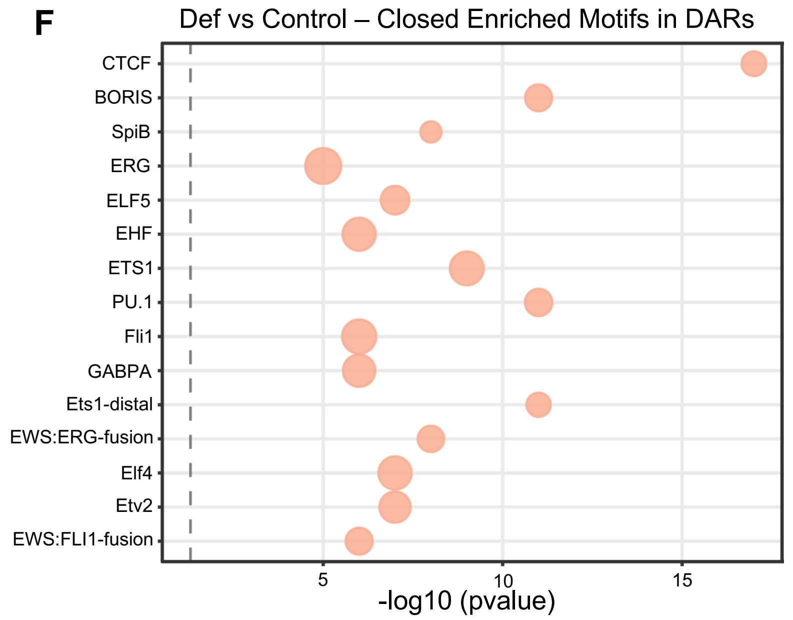


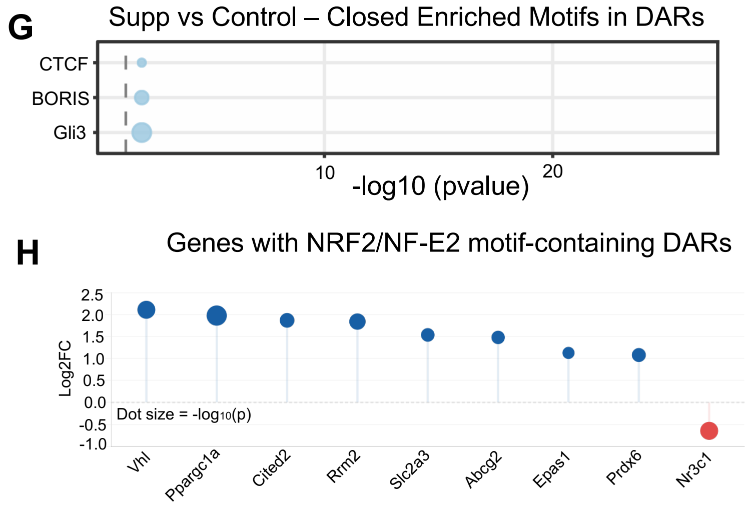


J

Self-renewal genes in Supp vs Control HSCs

| gene_name | pvalue | log2FoldChange |
|---|---|---|
| Sall4 | 0.04837137 | 1.001809063 |
| Bcl2 | 0.04278094 | 1.775057965 |

Polycomb-associated genes in Supp vs Control HSCs

I

| gene_name | pvalue | log2FoldChange |
|---|---|---|
| Sfmbt1 | 0.016051537 | 0.687281399 |
| Rybp | 0.030849301 | 0.590887433 |
| Cdkn2a | 0.003878128 | -1.610919345 |
| Cdkn2b | 0.045795957 | -1.641547986 |

K

Mitochondrial pyruvate handling genes in Supp vs Control HSCs

| gene_name | pvalue | log2FoldChange |
|---|---|---|
| Pdk4 | 0.0274163 | 2.624847966 |
| Hk2 | 0.01910286 | 2.290395571 |
